# Unraveling the Potential of Epicatechin Gallate from *Crataegus oxyacantha* in Targeting Aberrant Cardiac Ca2+ Signalling Proteins: An in-depth *In-Silico* Investigation for Heart Failure Therapy

**DOI:** 10.1101/2023.07.24.550267

**Authors:** J Praveen, M Anusuyadevi, KS Jayachandra

## Abstract

The cardiovascular sarcoplasmic reticulum (SR) calcium (Ca2+) ATPase is an imperative determinant of cardiac functionality. In addition, anomalies in Ca2+ handling protein and atypical energy metabolism are inherent in heart failure (HF). Moreover, Ca2+ overload in SR leads to mitochondrial matrix Ca2+ overload, which can trigger the generation of Reactive Oxygen Species (ROS), culminating in the triggering of the Permeability Transition Pore (PTP) and Cytochrome C release, resulting in apoptosis that leads to arrhythmias and numerous disorders. Although proteins involved in the molecular mechanism of Ca2+ dysfunction regarding mitochondrial dysfunction remains elusive, this study aims to assess the major Ca2+ handling proteins which may be involved in the Ca2+ malfunction that causes mitochondrial dysfunction and predicting the most effective drug by targeting the analyzed Ca2+ handling proteins through various *insilico* analyses.Thirteen proteins absorbed from interaction analysis were docked with four optimal phytochemicals from *Crataegus oxyacantha* (COC) to identify the potential agonist/antagonist against those anomalies causing Ca2+ handling signaling proteins. Furthermore,The ADMET profile of tyramine, vitexin, epicatechin, and epicatechin gallate was acclimated to evaluate potential drugability utilizing QikProp by Schrodinger.Keeping this in view, critical molecular docking evaluations were performed using Glide (Maestro), autodock, and autodock vina.Based on the results of 156 dockings by Maestro, auto-dock, and auto-dock vina, PKA Cα with epicatechin gallate exhibits good interaction. Therefore, a 2000ns molecular dynamics (MD) simulation was utilized to assess the feasible phytochemical epicatechin gallate - PKA Cα complex binding stability utilizing Desmond. As a result, the molecular dynamics simulation study confirmed that epicatechin gallate from COC has high possibilities to inhibit the aberrant cardiac Ca2+ signaling proteins due to its conformational rigidity.

## Introduction

The global leading cause of death is cardiovascular diseases (CVDs), an umbrella term including a variety of heart and blood vessel illnesses. Around 18 million fatalities worldwide occurred in 2019 due to these illnesses alone [1]. One of the major reasons for cardiac failure is Ca2+ overload in the SR of cardiomyocytes, which alters myocardial contractility, causing 80% of fatal arrhythmias with hypertrophic and dilated cardiomyopathies [2,3]. An individual, having an intensive myocardial infarction cause’s frequently progressing ischemia transpires discretionary to substantial atherosclerotic disease, results in anomalous signaling roles of Ca2+ contractile proteins, and is withal involved in the processes for apoptosis inhibition [4,5].

Post MI patients cannot survive more than 5 years due to arrythmias stunning because,during reperfusionwhen pH level is normalizedergo enhancement of Glucose (C₆H₁₂O₆) and oxygen (O2) cause aggregation of hydrogen ions over the plasma membrane of SR. Because of this, the concentration of hydrogen ions induces a sodium-hydrogen pump that stimulates sodium ions causing the anomalous function of changes in the sodium-Ca2+ pump which provokes ca2+ overload in the cytoplasm [6]. Fundamentally, an outrageous enhancement of intracellular Ca2+ leads to severe vicissitudes in cell signaling results perpetuated tachycardia and disintegration of cell function. These are eventually extrusion for impaired contraction and abnormal electrical activity [6,7]. For this reason, it is crucially essential to comprehend the mechanism behind the ca2+ overloading at the molecular level and distinguish the key proteins and transporters that contribute to the pathogenesis, to find the incipient strategies to treat coronary illness.

The general intracellular mechanism of cardiac Ca2+ begins its journey by Ryanodine Receptor 2 (RyR2) from SR release Ca2+ which suddenly increases the concentration level of cytosolic Ca2+ to binds with troponin C (TN-C) of actin/myosin to initiate muscle contraction. After that, Sarcoplasmic/Endoplasmic Reticulum Ca2+ ATPase (SERCA) recaptures the rapid removal of cytosolic Ca2+ return into the SR for initiating progress of the diastole [8,9]. Some Ca2+ also exits the cell through plasma membrane Ca2+ ATPase (PMCA) and sodium-Ca2+ exchanger channel (NCX) leading to the dissociation of Ca2+ from TN-C for cardiac muscle relaxation, and some Ca2+ is obtained by MCU from mitochondria for energy production for determining cell fate by preventing or triggering apoptosis [10,11]. To balance out the excitation-contraction coupling of the myocardium, Ca2+ balance needs to be maintained. In similar manner, assuming that there is an unevenness of Ca2+ inside the myocardium, there will be an unstable coupling is mentioned as arrhythmias and myocardial stunning.

Some proteins, such as NCX1 and ATP2A1/SERCA1, help to remove Ca2+ from the cytosol and replenish it back into the SR [12].Others, like CACNA1S, help to mediate Ca2+ influx during excitation-contraction coupling [13].S100 interacts with other proteins involved in Ca2+ handling pathways to help regulate cardiac function [14].However, an imbalance of Ca2+ in the heart can lead to cardiovascular diseases such as arrhythmias and myocardial stunning.Proteins such as BCL6 and PTEN can contribute to this imbalance by modulating Ca2+signaling [15].Additionally, proteins like COX2, BCL2, and P21 can also affect Ca2+ signaling and cell growth and apoptosis in the myocardium [16,17].Calreticulin is a protein located in the SR that plays a crucial role in regulating Ca2+ homeostasis by regulating the expression of other proteins involved in Ca2+ handling pathways [18].Hence,the interaction of various proteins is necessary for maintaining proper Ca2+ balance in the heart and preventing cardiovascular diseases. Therefore, it is crucial to maintain the delicate balance of Ca2+ in the heart to ensure proper cardiac functioning.

Nowadays, natural products meet a paramount and clamant role in discovering efficacious drugs to treat critical diseases[19]. There is a developing assemblage of evidence that *COC* Linn. (Rosaceae) commonly known as Hawthorn extracts exhibit cardioprotective properties in both in-vivo and in-vitro studies[20]. A primary purpose of the present study is to determine the binding efficiency of major secondary metabolites of *COC* with Ca2+ overload-induced Myocardial Infarction-associated proteins through insillico studies.

## Materials and methods

### Selection of target proteins

Proteins that play an important role in inducing Ca2+ overload have been data mined from string database (https://string-db.org/), based on various literature reviews and interaction analyses. Thusly, thirteen proteins were considered as a target for molecular docking analysis through glide (Maestro), autodock, autodock vina. The protein sequence was retrieved from PDB (https://www.rcsb.org/pdb/) and Uniprot (https://www.uniprot.com).

### Selection of phytochemicals

The widely used cardiotonic herb COC (Hawthorn) is usually acquainted to treat various cardiovasculardiseases^20^.A literature search yielded four important phytochemical substances Vitexin, Tyramine, Epicatechin, and Epicatechin Gallate of COC (COC), which are employed as primary therapeutic components against proteins that induce aberrant molecular expression in cell^22,23^.These compounds structures were obtained from the PubChem Compound database (https://pubchem.ncbi.nlm.nih.gov/).

### Active site prediction

The information about the active site region was obtained from PDB database records from Discovery studio Visualizer. Several proteins do not have active site information in the PDB. Anticipating active site wasn’t difficult to predict. However, determining the optimal active site was more difficult. Blind docking is a sophisticated technique that analyses the whole surface of the protein target for potential ligand-binding sites [24]. In blind docking, the whole surface of a protein is docked to recognize potential interactions between protein and its ligand and to distinguish the most probable restricting mode for that protein [25,26,27]. CB-Dock (http://cao.labshare.cn/cb-dock/,), which reveals protein-binding regions with coordinates relying upon the autodock vina [28], was utilized for the blind docking.

### Protein preparation

The protein structure retrieved from the PDB database couldn’t be utilized directly for docking analyses due to the existence of multiple protein chains, water molecules, cleansers, little atoms, co-elements, and metal molecules. After crucial processing’s, unnecessary molecules and protein chains were eliminated. A legitimate outcome requires adding polar hydrogen atoms to electronegative atoms and tracking down the missing atoms in the residue. From that point onward, the kollmans charges and non-bonded atoms were figured.

### Ligand preparation

Pymol was utilized since Autodock and Autodock vina both support ligands in PDB format. In this case, the ligand molecule should compute the charges without any water molecule. In addition, the maximum number of possible torsions was determined subsequently to check the number of rotatable bonds. After that, the ligand was generated, and the docking analysis was performed using the optimized ligands.

### Protein-ligand interaction using Autodock

Autodock vina is a docking program based on the Lamarckian genetic algorithm search method with an empirical free energy force field [29,30]. Simultaneously, Autodock tools (graphical UI) have been developed for various working frameworks and were utilized for constructing auto grid points as well as imagining docked protein-ligand complexes. After that, A target site and grid box with a dispersing of 0.375 Å were made on the protein structure utilizing information from the PDB records and blind docking. Autodock 4.2.6 was utilized to test several docking parameter options including total energy evaluations per docking run and the total number of dockings runs to determine the optimal docking conditions between protein and ligands. During the docking, a maximum of 2.5 million energy assessments with 30 genetic algorithm runs were culled. As a consequence, thirteenprotein-ligand docking analysis were conducted with the maximum number of energy evaluations for the precise docking result with ideal positioning. All other docking parameters were left at the default values and customarily, the docking run has culminated when the number of energy evaluations is reached.

### Protein-ligand interaction using Autodock vina

Autodock vina provides a viable solution for protein-ligand docking based on the quick gradient-optimization conformational search and a simple scoring function algorithm [31]. Besides, protein-ligand docking was culminated by utilizing autodock vina to evaluate the hydrogen bond interactions and binding affinities to prognosticate results more preciously. Following the minimization process, the grid box resolution was computed based on the PDB record reviewed by the discovery studio visualizer. Through the blind docking by utilizing CB-Dock (http://clab.labshare.cn/cb-dock/), the coordinates of proteins with obscure PDB records were inferred. On the other hand, the grid dimensions were set as default. In this manner, five phytochemicals were docked independently with thirteen proteins in order to obtained the best protein for further study.

### Protein ligand Docking by Maestro

In this study, protein-ligand docking was performed using the GLIDE by Schrodinger to predict the binding mode and binding energy of thirteen proteins with the secondary metabolites of COC. The protein structure was prepared by removing waters and cofactors, ensuring that the structure was cleaned and minimized using PDB2PQR and PROPKA. The ligand structure was obtained in SDF format in PubChem, and was prepared by ensuring that it was in a minimized, cleaned format with all explicit hydrogens added. After the uploading of protein and ligand, the docking simulation was performed using GLIDE to evaluate the binding energy and visualized using Discovery studio visualizer.

### Molecular dynamics simulation of PKA Cα – epicatechin gallate complex

Desmond was utilized to examine the conformational changes in the PKA Cα – epicatechin gallate complex. The PKA Cα – epicatechin gallate complex was placed in an orthorhombic cubic box for dynamics simulations, TIP3P dihydrogen monoxide molecules and buffers were filled amongst the protein and box’s edge. In addition, to neutralize the system, the volume of the complex type boundary condition has been computed using counterions. For the purpose of the energy calculation, the default temperatures of 300 K and 1.01325 bar were maintained by means of the Nose-Hoover thermostat and the Martyna Tobias-Klein Barostat algorithm. Besides NPT ensembles simulations were conducted for 2000ns.

## Result

A string (https://string-db.org/) database analysis revealed major thirteen proteins that are associated with causing Ca2+ imbalance. As a result, an intensive docking and dynamics investigation was carried out utilizing several software and servers to find optimal phytochemicals of COC prospective drugs against proteins embroiled in Ca2+ overload.

### Proteins implicated in the signaling of Ca2+ overloading were retrieved with the precious crystal structure

In the drug discovery process, it is fundamental to survey the activity of a major thirteen protein as a target by determining its crystal structure retrieved from the Protein Data Bank (https://www.rcsb.org/) as illustrated in table 1.

**Table 1:** Important proteins involved in the progression of pre-Myocardial Infarction in ca2+ overloading.

| S. No | Protein<sup>a</sup> | Protein Function<sup>b</sup> | Uniprot ID<sup>c</sup> | PDB ID<sup>d</sup> |
| --- | --- | --- | --- | --- |
| 1 | NCX1 | regulation of cytoplasmic Ca<sup>2+</sup> levels | P23685 | 2DPK |
| 2 | ATP2A1/SERCA1 | responsible for the reuptake of cytosolic Ca<sup>2+</sup> | P04191 | 6HEF |
| 3 | CACNA1S | Calmodulin mediator | P0DP23 | 2VAY |
| 4 | S100 | contributing to cellular Ca<sup>2+</sup> signaling | P30801 | 1A03 |
| 5 | Ca<sup>2+</sup>/calmodulin-dependent protein kinase II | involved in SR Ca<sup>2+</sup> transport | Q13555 | 2UX0 |
| 6 | PKA C-α (Protein kinase A catalytic subunit) | Involved in process of Phosphorylates | P17612 | 2GU8 |
| 7 | Triadin | Membrane-bound Ca<sup>2+</sup>-sensing protein | Q12797 | 6QA5 |
| 8 | bcl6 | Transcriptional repressor | P41182 | 5N20 |
| 9 | PTEN | protein phosphatase and dephosphorylating | P60484 | 1D5R |
| 10 | COX2 | cyclooxygenase and peroxidase in the biosynthesis pathway | P35354 | 5F1A |
| 11 | BCL2 | Suppresses apoptosis | P10415 | 1G5M |
| 12 | P21 | activation of Ras protein signal transduction | P01112 | 5P21 |
| 13 | CALRETICULIN | Ca<sup>2+</sup> -binding chaperone | P27797 | 3POW |

<sup>a</sup>a name of thirteen proteins as a target by determining its crystal structure retrieved from the Protein Data Bank,
<sup>b</sup>A molecular and biological functions of protein. <sup>c</sup>Uniprot ID for the targeted proteins, <sup>d</sup>PDB ID for the targeted proteins

### Finalist bioactive compounds were retrieved from cardio therapeutic herb COC

In antiquated times, COC (Hawthorn) have been utilized as therapeutic plants are attributed to their antioxidant constituents [21].For these ongoing studies, epicatechin, epicatechin gallate, vitexin, and tyramine, as well as their structures were accumulated from previous literature investigations [22,23]. The verified structures were retrieved from PubChem database (https://pubchem.ncbi.nlm.nih.gov/) as illustrated in Figure (1).

### An insillico approach to quality control of drugs based on ADME prediction

In the development of drug candidates, unfavourable absorption, distribution, metabolism, and elimination (ADME) characteristics have been cited as a vital factor in forestalling success^32^. An assortment of potential SASA properties, octanol/water partition coefficient, Predicted Blood-Brain partition coefficient, aqueous solubility, and human oral absorption were assessed with QikProp (Schrodinger).

### An exhaustive investigation of binding affinities of finalist bioactive compounds of COC with proteins involved in Ca2+ overload signalling utilizing Autodock Vina

According the literature studies, it was advised to test both autodock / autodock vina wisely for specific systems regardless of their similar protein compositions [33]. Hence, this study compares the values of both binding energy of autodock and binding affinity of autodock vina by examining novel inhibitors for ca2+ overloading in myocardium. In present study, binding affinity oriented molecular docking simulations were performed on thirteen proteins with four secondary metabolites of COC.

A better binding affinity was obtained epicatechin gallate with P21, PKA C-α, COX2 and Vitexin with BCL2, P21(Table 4). The 3D and 2D structure of P21, PKA C-α, COX2 and Vitexin with BCL2, P21 docking results was as follow as

### A comprehensive binding energy investigation of molecular docking of Finalist bioactive compounds with Proteins embroiled in the signaling of Ca2+ overloading utilizing Autodock

The molecular docking of thirteen proteins with selected potential secondary metabolites were docked with 2.5 million energy assessments with thirty genetic algorithms were culled with maximum number of energy evaluations for the precise docking results with ideal positioning.

### A comprehensive study was undertaken to determine the binding energy and glide energy between finalist bioactive compounds of COC and proteins involved in Ca2+ overload signaling by utilizing maestro: -

In our study, we used the Glide software from Schrodinger to perform protein-ligand docking on a thirteen targeted protein of interest (Table 1) with four ligands (Figure 1). The results of the docking simulations showed that several of the ligands had high scoring binding poses, with the glide scores in the range of −5.0 kcal/mol to −8.0 kcal/mol. These scores indicate a strong predicted binding affinity of the ligands to the protein. The results of the binding poses were indicated (Table 7,8,13,14,15) and the top three results were also indicated with 3d and 2d structures. As the Glide results, the epicatechin gallate with 2GU8 has high accuracy and able to be a potential lead compound for further drug development and validation with experiments.

**Figure 1:**
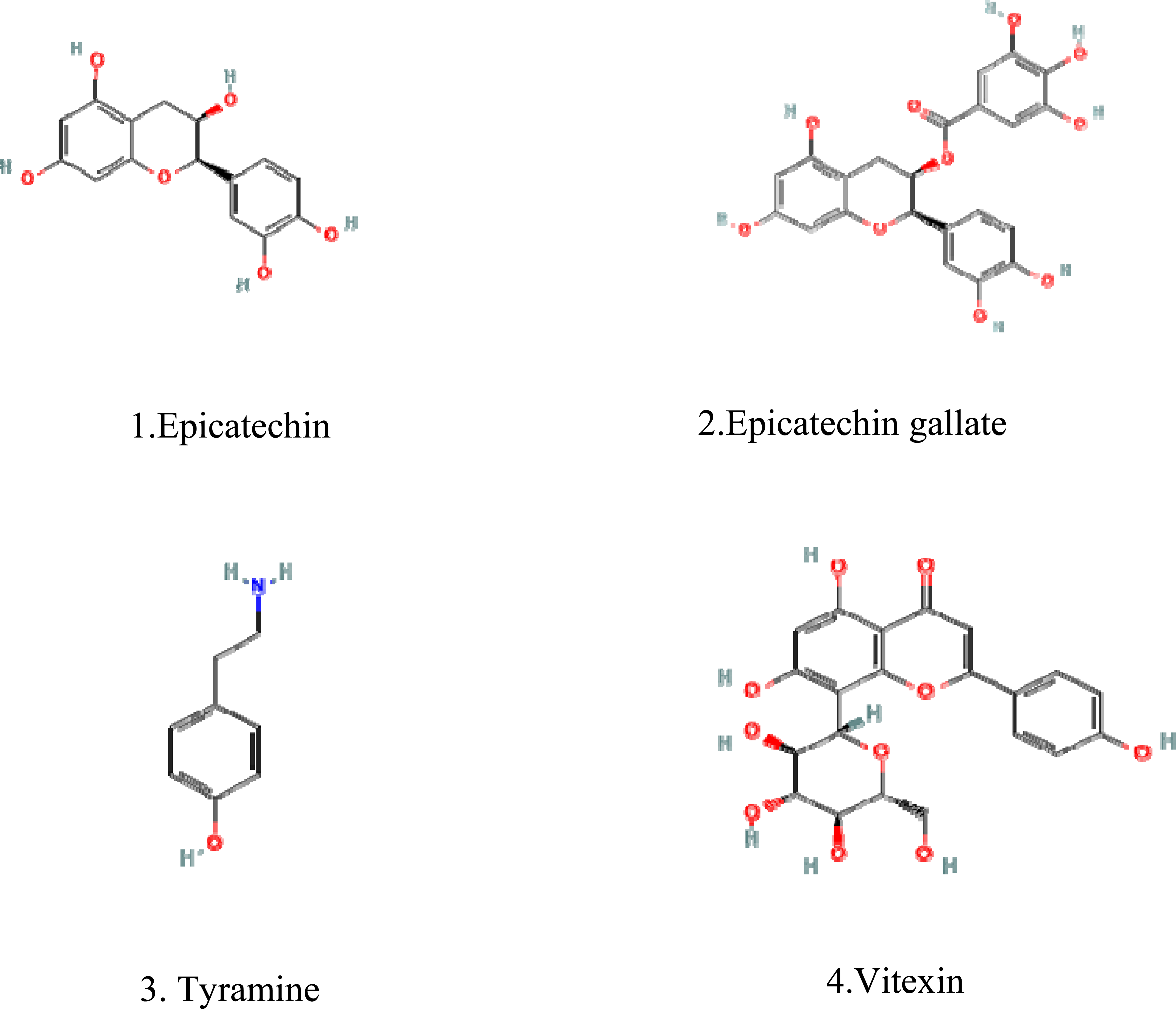
The structure of the four significantly active secondary metabolites of *COC* obtained from PubChem.

### An in-depth examination of the relationship between ligand binding and targeted protein underlying changes is carried out by utilizing molecular dynamics simulation studies

As indicated by the molecular docking studies, 26 best interactions have been identified between protein and inhibitor based on Glide (Maestro), Autodock and Autodock vina data, and the compound epicatechin gallate was additionally assessed for conceivable atomic subtleties. Hence, Molecular dynamics simulations were directed by utilizing Desmond software to assess the stability of PKA Cα with epicatechin gallate. As a result, 2000ns simulation was investigated to determine the stability of the protein-inhibitor complex by considering H-bond interactions, RMSD and energy value. Additionally, the figure (4,8) incorporates the histogram chart and two-dimensional interaction poses.

**Figure 2:**
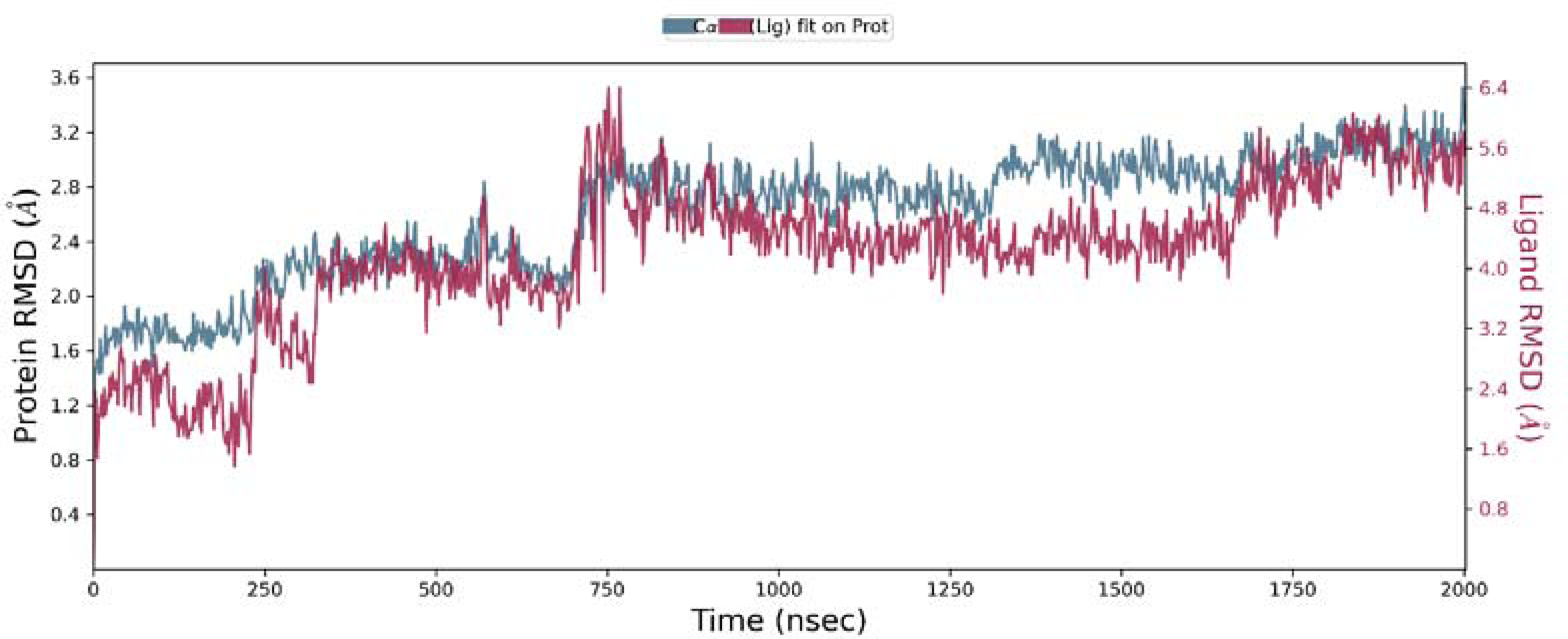
Root Means Square Deviation (RMSD) plot for PKA C-α – Epicatechin gallate during 2000ns of molecular dynamics simulation. Protein PKA C-α is shown in the blue colour and Epicatechin gallate in the red colour.

## Discussion

By assessing the significant Ca2+ handling proteins engaged with the pathophysiological functions by ca2+ causing mitochondrial dysfunction, this study aims to anticipate the most effective secondary metabolites from COC targeting on the major anomalies signaling of ca2+ handling proteins in cardiomyocytes. In addition to better comprehension of cardiac diseases at the molecular level, identification of impacted key proteins and transporters engaged with pathogenesis has prompted to the novel therapeutic methodologies.

Ca2+ plays a several signalling roles including muscle contraction and relaxation in cardio vascular system [32]. In cardiomyocytes, the systole and diastole represent rapid variations in intracellular Ca2+ contraction, which is governed by excitation – contraction coupling. Through the voltage gated L-type Ca2+ channel, extracellular Ca2+ is entered into the cytosol during systole following an action potential depolarizing the sarcolemma of the plasma membrane. Afterwards, Ca2+ is largely effluxes into the cytosol in the wake of binding to ryanodine receptor (RyR2) and ca2+ in the cytosol also enacts ATP subordinate contractile proteins for muscle contractions. For muscle relaxation, around about a third of Ca2+ is withdrawn from the cytosol by various proteins, while remaining is reabsorbed into endoplasmic reticulum by the cardiac SR Ca2+ ATPase (SERCA2a) [34]. Dysfunction of Ca2+ transport and Ca2+ handling proteins leads to defects in EC coupling resulting in diminished contractility and cardiac output cause Ca2+ spill in endoplasmic reticulum.

In various animal models, flavonoids and procyanidins have been displayed to exert similar effects on contractility as those found in the leaves and fruits of COC [35]. Hence, the secondary metabolites in the COC extract were identified in the literature and their structures were retrieved from the PubChem database. A physiochemical description of Epicatechin, epicatechin gallate, tyramine and vitexin, as well as ADME parameters, pharmacokinetics, and medicinal chemistry friendliness were anticipated by QikProp in table 2 and the outcomes were in the suggested range.

**Table 2:** ADME prediction of Epicatechin, Epicatechin Gallate, Tyramine and Vitexin were assessed with QikProp (Schrodinger)

| Compound Name | MW<sup>a</sup> | Donors HB<sup>b</sup> | Acceptors HB<sup>c</sup> | SASA<sup>d</sup> | QPlogPo/w<sup>e</sup> | QPlogBB<sup>f</sup> | QPlogS<sup>g</sup> | %Human oral Absorption<sup>h</sup> |
| --- | --- | --- | --- | --- | --- | --- | --- | --- |
| Epicatechin | 442.378 | 1 | 2.25 | 683.541 | 2.898 | -3.604 | -6.78 | 51.548 |
| Epicatechin Gallate | 290.272 | 5 | 5.45 | 509.365 | 0.491 | -1.848 | -2.586 | 60.951 |
| Tyramine | 137.181 | 4 | 0.75 | 326.491 | 0.147 | -0.447 | -0.702 | 80.743 |
| Vitexin | 432.383 | 5 | 9.5 | 517.081 | -1.323 | -2.006 | -2.815 | 35.411 |

<sup>a</sup> Molecular weight (acceptable range: < 500), <sup>b</sup> Hydrogen bond donor (acceptable range: <500), <sup>c</sup>Hydrogen bond acceptor (acceptable range:<10), <sup>d</sup> Total solvent accessible surface area using a probe with a 1.4 radius (acceptable range: 300-1000), <sup>e</sup> predicted octanol/water partition coefficient (acceptable range: -2 - 6.5), <sup>f</sup> Predicted Blood-Brain partition coefficient (acceptable range: -3 – 1.2), <sup>g</sup>predicted aqueous solubility, S in mol/dm-3 (acceptable range: -6.5 -0.5), <sup>h</sup> predicted human oral absorption on 0 to 100% scale (<25% is poor and 80% is high)

**Table 3:**
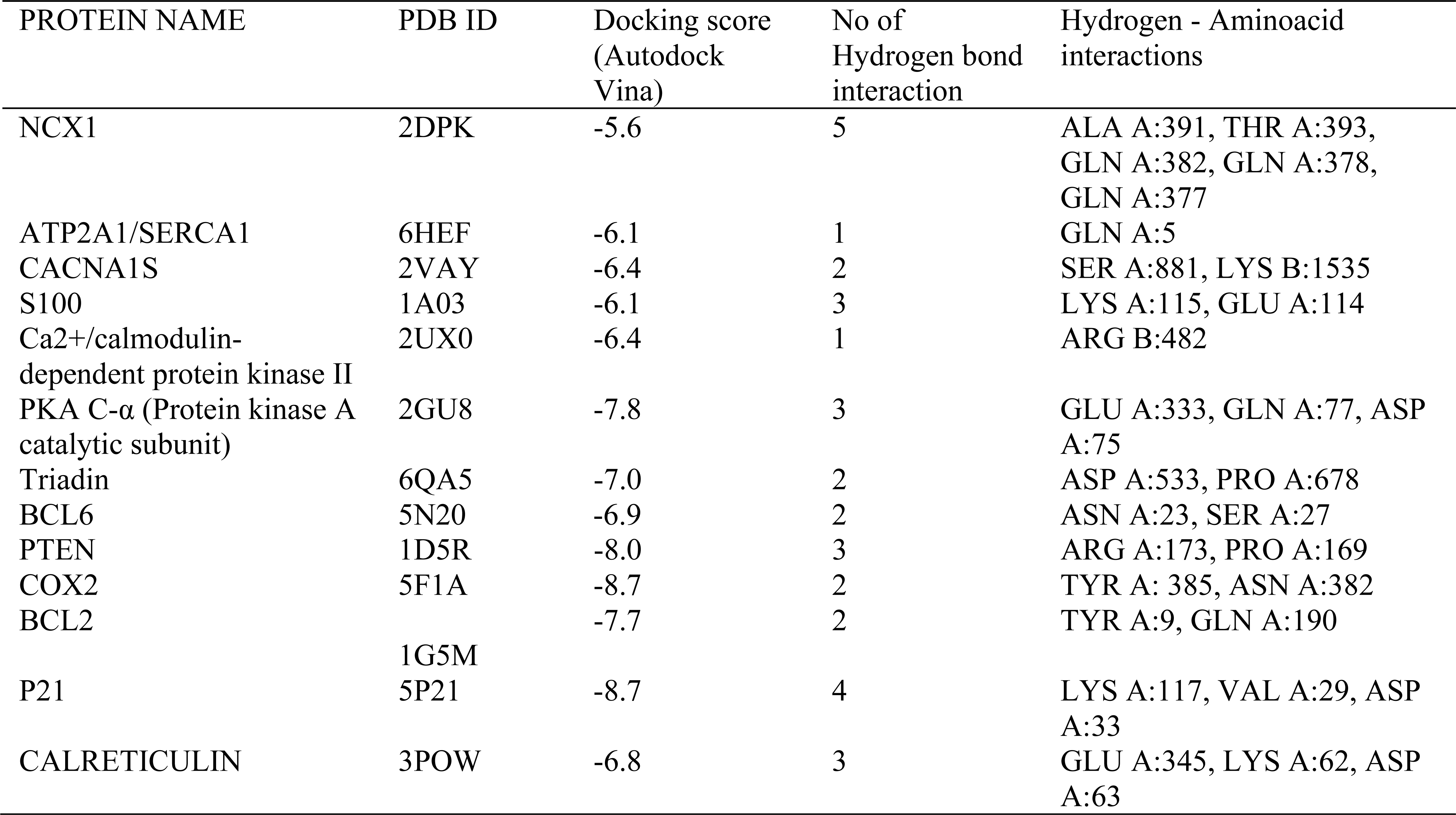
A comprehensive binding affinity results with hydrogen bonding interactions of molecular docking of Epicatechin with Proteins embroiled in the signaling of Ca2+ overloading through Autodock vina.

**Table 4:** A comprehensive binding affinity results with hydrogen bonding interactions of molecular docking of Epicatechin gallate with Proteins embroiled in the signaling of Ca2+ overloading through Autodock vina.

| PROTEIN NAME | PDB ID | Docking score<br>(Autodock<br>Vina) | No of Hydrogen<br>bond interaction | Hydrogen - Aminoacid<br>interactions |
| --- | --- | --- | --- | --- |
| NCX1 | 2DPK | -6.5 | 6 | THR A:380, GLN A:378,<br>THR A:393, GLU A:439,<br>THR A:389, GLN A:382 |
| ATP2A1/SERCA1 | 6HEF | -6.3 | 2 | ARG A:924, PHE A:809 |
| CACNA1S | 2VAY | -6.9 | 4 | MET A:145, SER A:81,<br>ALA A:147, LYS B:1535 |
| S100 | 1A03 | -6.5 | 5 | LEU A:112, GLU A:14<br>GLN B:1530, ARG B: 1534 |
| Ca2+/calmodulin-<br>dependent protein kinase II | 2UX0 | -8.1 | 1 | SER B:496 |
| PKA C-α (Protein kinase A<br>catalytic subunit) | 2GU8 | -9.0 | 2 | TYR A:330, GLY A:55 |
| Triadin | 6QA5 | -7.9 | 6 | ASN A:618, THR A:680,<br>PRO A:678, ASP A:553 |
| BCL6 | 5N20 | -7.2 | 4 | GLN A:42, ASN A:84, ASN<br>A:23 |
| PTEN | 1D5R | -8.0 | 4 | LEU A:152, LYS A:332,<br>ASN A:329, ASN A:323 |
| COX2 | 5F1A | -8.6 | 4 | TYR A:385, HIS A:214,<br>HIS A:207, PHE A:210 |
| BCL2 | 1G5M | -8.2 | 5 | GLY A:193, TRP A:195,<br>TYR A:9, ASN A:11, ASN<br>A:182 |
| P21 | 5P21 | -9.5 | 4 | LYS A:117, ASP A:119,<br>LYS A:147, ALA A:146 |
| CALRETICULIN | 3POW | -8.1 | 4 | GLU A:345, ASN A:329,<br>THR A:350 |

**Table 5:** A comprehensive binding affinity results with hydrogen bonding interactions of molecular docking of Tyramine with Proteins embroiled in the signaling of Ca2+ overloading through Autodock vina.

| PROTEIN NAME | PDB ID | Docking score<br>(Autodock Vina) | No of Hydrogen<br>bond interaction | Hydrogen - Aminoacid<br>interactions |
| --- | --- | --- | --- | --- |
| NCX1 | 2DPK | -3.9 | 1 | ALA A:391 |
| ATP2A1/SERCA1 | 6HEF | -4.5 | 2 | GLN A:56, PRO A:312 |
| CACNA1S | 2VAY | -5.0 | 1 | ASP A:78 |
| S100 | 1A03 | -4.1 | 2 | MET A:145 |
| Ca2+/calmodulin-dependent<br>protein kinase II | 2UX0 | -4.5 | 1 | ASN A:411 |
| PKA C-α (Protein kinase A<br>catalytic subunit) | 2GU8 | -5.6 | 3 | SER A:553, ASP A:184,<br>LYS A:72 |
| Triadin | 6QA5 | -5.3 | 1 | HIS A:555 |
| BCL6 | 5N20 | -4.3 | 0 | NIL |
| PTEN | 1D5R | -5.5 | 1 | ALA A:328 |
| COX2 | 5F1A | -6.4 | 2 | GLN A:203, TYR A:385 |
| BCL2 | 1G5M | -4.4 | 1 | ALA A:131 |
| P21 | 5P21 | -6.1 | 5 | ASP A:30, ASP A: 119,<br>LYS A:147, ALA A:146,<br>VAL A:29 |
| CALRETICULIN | 3POW | -5.3 | 3 | GLU A: 345, ASP A:63,<br>LYS A:62 |

**Table 6:** A comprehensive binding affinity results with hydrogen bonding interactions of molecular docking of Vitexin with Proteins embroiled in the signaling of Ca2+ overloading through Autodock vina.

| PROTEIN NAME | PDB ID | Docking score (Autodock Vina) | No of Hydrogen bond interaction | Hydrogen - Aminoacid interactions |
| --- | --- | --- | --- | --- |
| NCX1 | 2DPK | -5.8 | 2 | THR A:493, ASN A:455 |
| ATP2A1/SERCA1 | 6HEF | -7.4 | 5 | PRO A:312, ASP A:254, ASP A:59, THR A:316, GLN A:56 |
| CACNA1S | 2VAY | -7.3 | 5 | SER A:81, LYS B:1535, THR A:79, MET A:76 |
| S100 | 1A03 | -6.6 | 0 | NIL |
| Ca<sup>2+</sup>/calmodulin-dependent protein kinase II | 2UX0 | -7.5 | 1 | ARG A:482 |
| PKA C-α (Protein kinase A catalytic subunit) | 2GU8 | -8.1 | 3 | HIS A:68, TYR A:122, GLU A:176 |
| Triadin | 6QA5 | -7.5 | 5 | ASN A:458, SER A:427, ASP A:428, GLN A:431 |
| BCL6 | 5N20 | -6.9 | 2 | HIS A:116, SER A:93 |
| PTEN | 1D5R | -7.4 | 4 | ARG A:159, GLN A:171, ASP A:92 |
| COX2 | 5F1A | -9.3 | 1 | THR A:206 |
| BCL2 | 1G5M | -8.2 | 2 | TRP A:195, GLN A:190 |
| P21 | 5P21 | -8.9 | 3 | LYS A:147, ASP A:33, SER A:17 |
| CALRETICULIN | 3POW | -7.5 | 7 | GLN A:26, PHE A:27, LEU A:28, ASP A:29, SER A:80 |

**Table 7:**
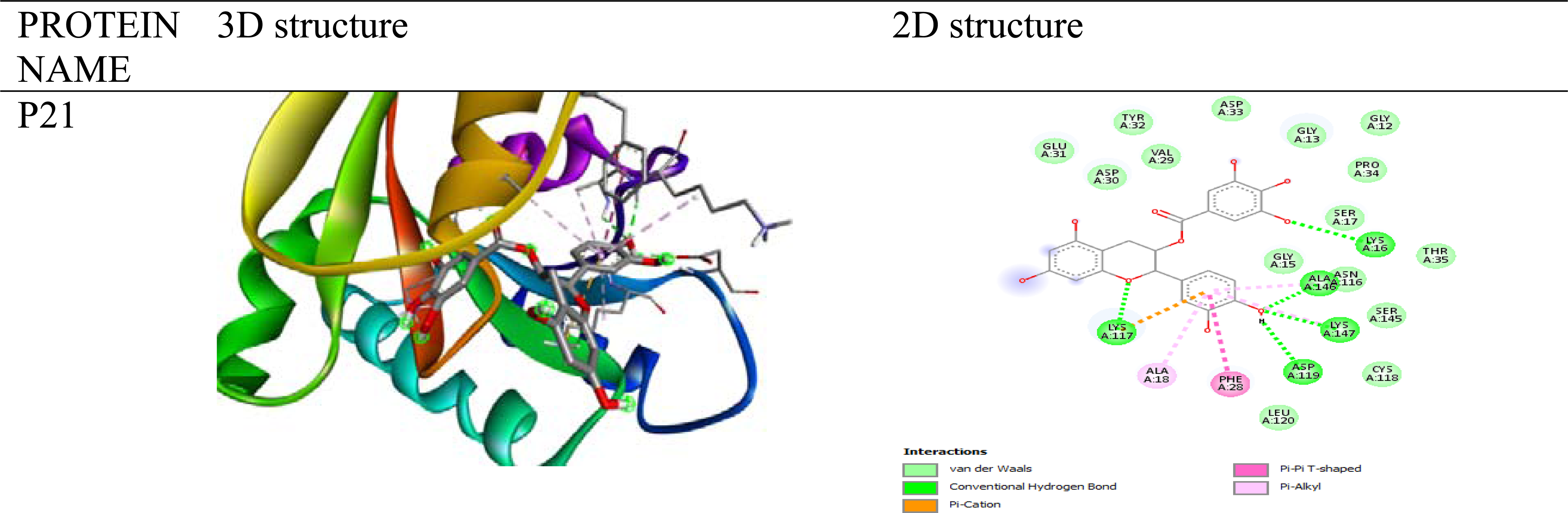
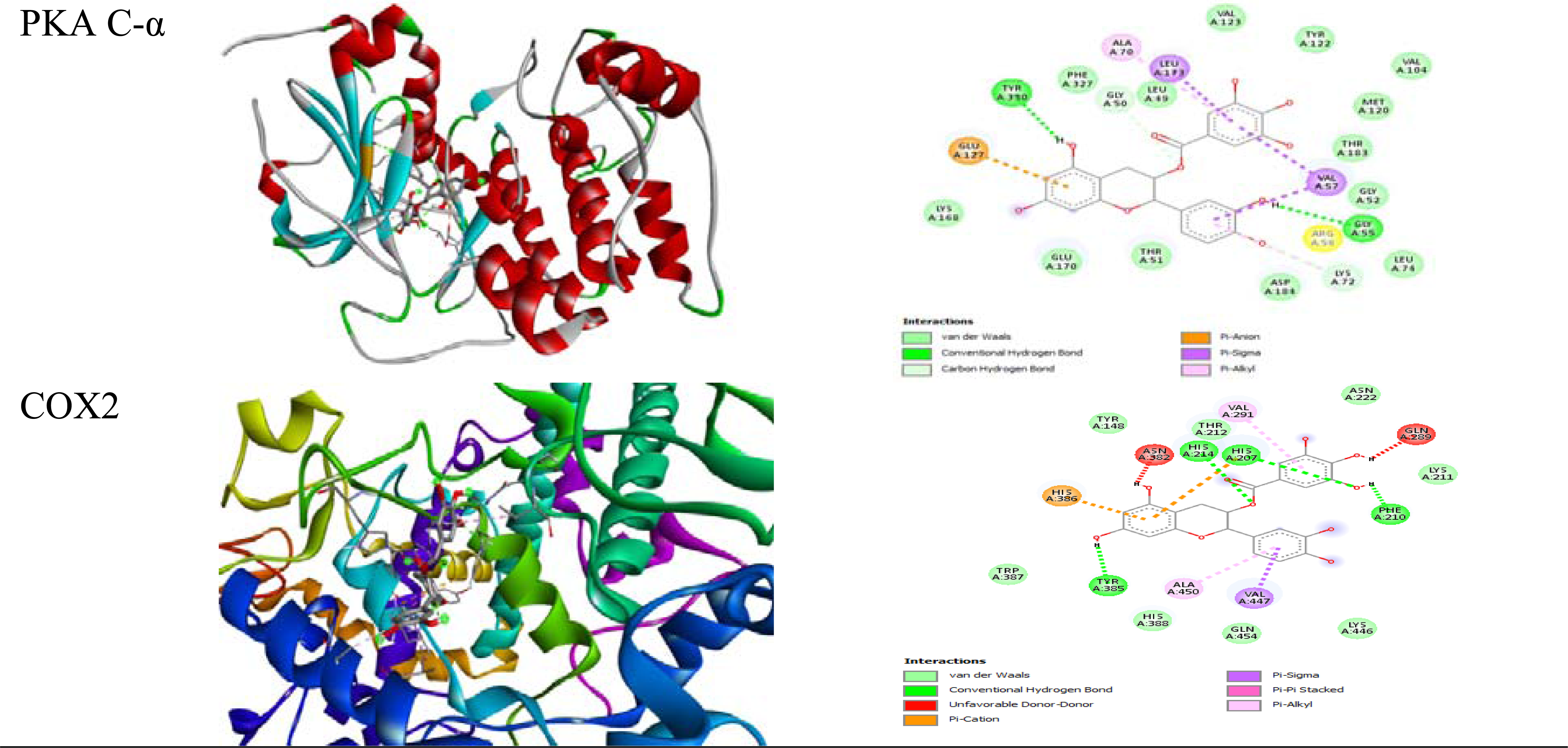
3D and 2D structure of Epicatechin gallate with P21, PKA C-α, COX2.

**Table 8:**
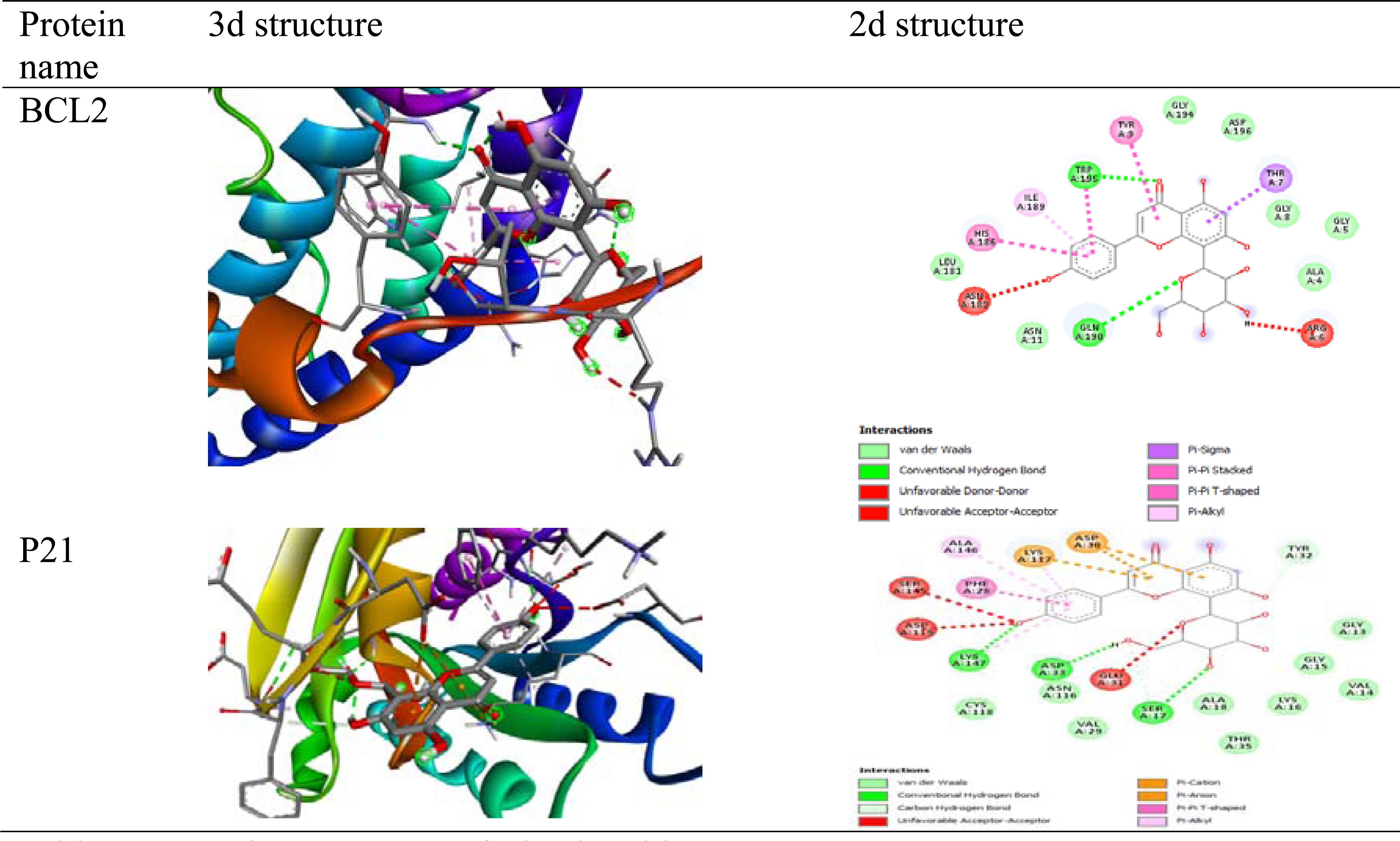
3D and 2D structure of Vitexin with P21, PKA C-α, COX2.

**Table 9:** A comprehensive binding energy results with hydrogen bonding interactions of molecular docking of Epicatechin with Proteins embroiled in the signaling of Ca2+ overloading through Autodock.

| PROTEIN NAME | PDB ID | Docking score<br>(Autodock) | No of<br>Hydrogen<br>bond<br>interaction | Hydrogen - Aminoacid<br>interactions |
| --- | --- | --- | --- | --- |
| NCX1 | 2DPK | -3.54 | 5 | ASP A:421, GLU A :451,<br>GLU A: 452, ASP A: 447 |
| ATP2A1/SERCA1 | 6HEF | -4.31 | 2 | LEU A: 797 |
| CACNA1S | 2VAY | -3.26 | 4 | MET A:51, LYS A: 75,<br>LYS B: 1522 and GLU<br>A:84 |
| S100 | 1A03 | -2.08 | 2 | GLN A:71, ARG A: 62 |
| Ca2+/calmodulin-<br>dependent protein kinase II | 2UX0 | -6.41 | 5 | GLU A: 432, GLU A:<br>430, ARG A: 502, TYR<br>A: 418 |
| PKA C-α (Protein kinase A<br>catalytic subunit) | 2GU8 | -7.20 | 5 | VAL A:123, GLU A:121,<br>THR A:183, LYS A:72,<br>ASP A:184 |
| Triadin | 6QA5 | -4.11 | 3 | ARG A:526, HIS A: 493,<br>GLY A:521 |
| BCL6 | 5N20 | -3.49 | 4 | ALA A:15, MET A: 90,<br>ALA A:52, CYS A:53 |
| PTEN | 1D5R | -3.38 | 4 | ARG A:130, ASP A:92,<br>GLN A :171 |
| COX2 | 5F1A | -8.01 | 5 | TYR A: 385, THR A:<br>206, ALA A: 202, TRP<br>A:387, ASN A: 382 |
| BCL2 | 1G5M | -4.94 | 5 | GLY A:193, GLN A:190,<br>TYR A:9, ASN A:11,<br>ASN A:182 |
| P21 | 5P21 | -6.02 | 7 | GLU A:31, ASP A: 33,<br>SER A:17, ALA A:11,<br>GLY A:15, LYS A :16 |
| CALRETICULIN | 3POW | -6.28 | 6 | TYR A:338, GLU A:345,<br>THR A:350, ASP A: 63,<br>GLN A:26 |

**Table 10:**
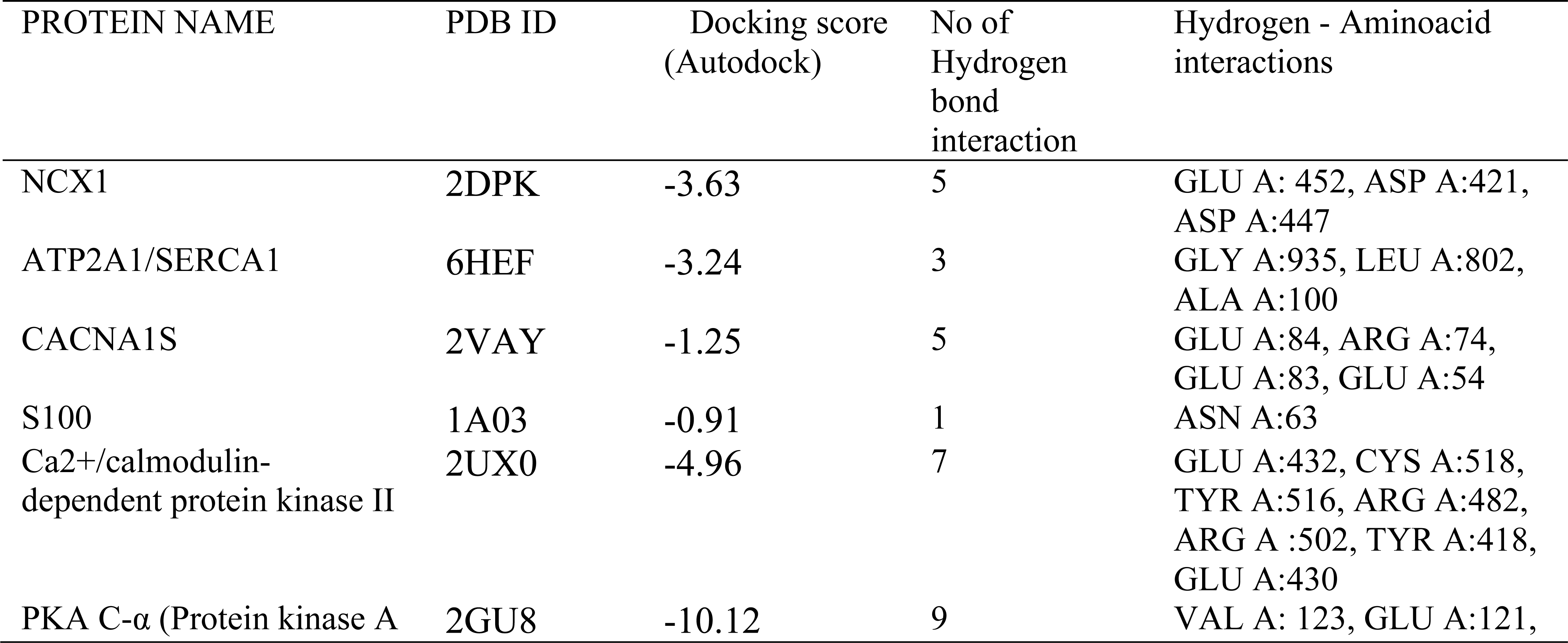
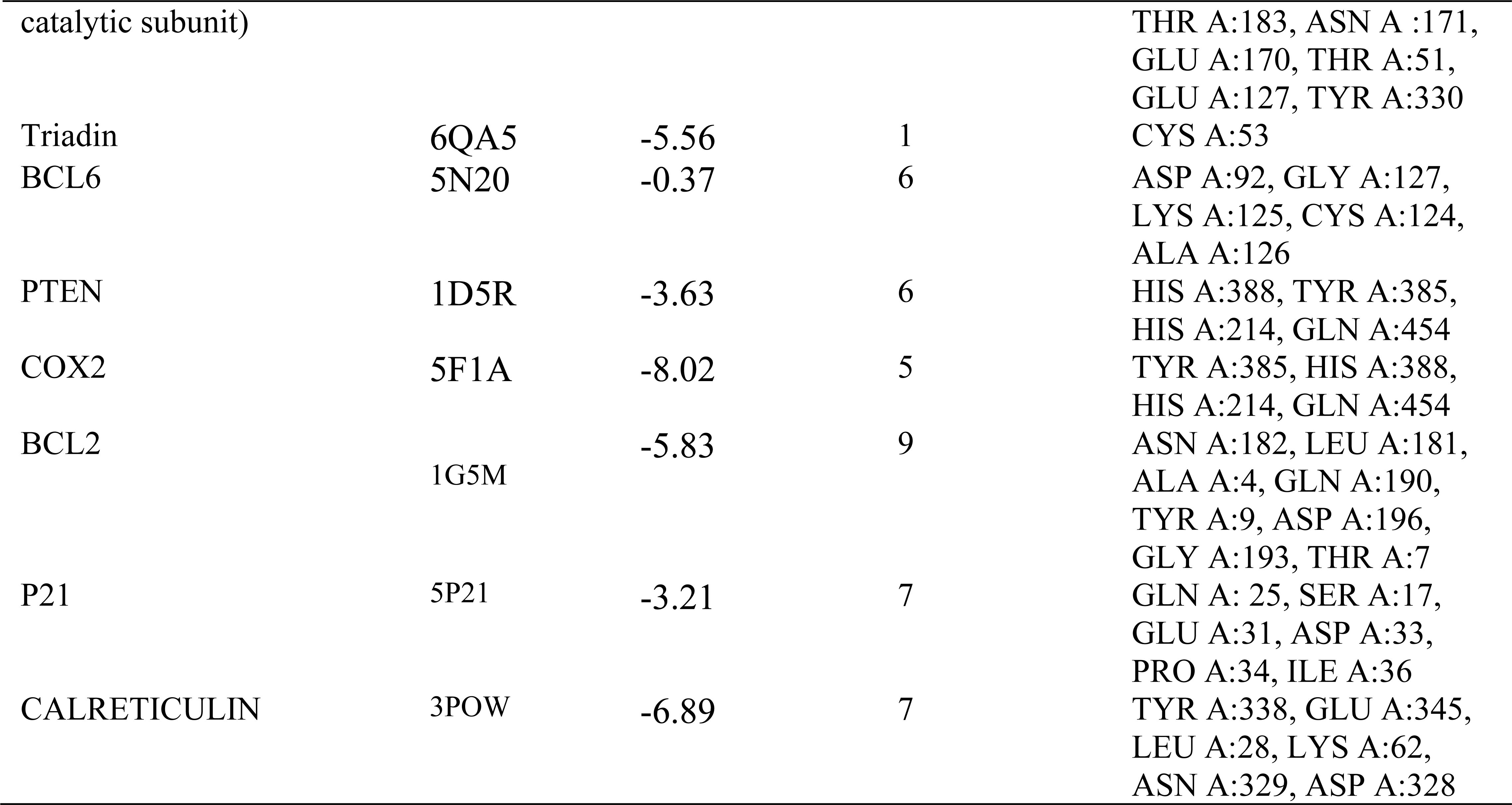
A comprehensive binding energy results with hydrogen bonding interactions of molecular docking of Epicatechin Gallate with Proteins embroiled in the signaling of Ca2+ overloading through Autodock.

**Table 11:** A comprehensive binding energy results with hydrogen bonding interactions of molecular docking of tyramine with Proteins embroiled in the signaling of Ca2+ overloading through Autodock.

| PROTEIN NAME | PDB ID | Docking score<br>(Autodock) | No of Hydrogen<br>bond interaction | Hydrogen -<br>Aminoacid<br>interactions |
| --- | --- | --- | --- | --- |
| NCX1 | 2DPK | -7.06 | 5 | GLU A:454, GLU<br>A:451, ASP A:447,<br>ASP A:421 |
| ATP2A1/SERCA1 | 6HEF | -3.85 | 2 | SER A:936, SER<br>A:940 |
| CACNA1S | 2VAY | -4.99 | 3 | GLU A: 54, TYR<br>B:1524 |
| S100 | 1A03 | -3.13 | 3 | GLU A: 67 |
| Ca<sup>2+</sup>/calmodulin-<br>dependent protein kinase II | 2UX0 | -6.59 | 4 | GLU A:430, TYR<br>A:418, ARG A:502 |
| PKA C-α (Protein kinase A<br>catalytic subunit) | 2GU8 | -5.35 | 3 | GLU A: 127, LEU<br>A:49, VAL A:123 |
| Triadin | 6QA5 | -4.53 | 3 | ASP A:487, ASP<br>A:459 |
| BCL6 | 5N20 | -2.75 | 4 | MET A:90, CYS<br>A:53, ALA A:52 |
| PTEN | 1D5R | -1.69 | 4 | ASP A:92, ARG<br>A:130, CYS A:124 |
| COX2 | 5F1A | -5.72 | 3 | TYR A:385, THR<br>A:206, TRP A:387 |
| BCL2 | 1G5M | -5.15 | 2 | ASP A:111, SER<br>A:105 |
| P21 | 5P21 | -5.15 | 3 | THR A:35, ASP<br>A:38, GLU A:31 |
| CALRETICULIN | 3POW | -7.45 | 4 | ASP A:63, THR<br>A:350, LYS A:62 |

**Table 12:** A comprehensive binding energy results with hydrogen bonding interactions of molecular docking of Vitexin with Proteins embroiled in the signaling of Ca2+ overloading through Autodock.

| PROTEIN NAME | PDB ID | Docking score<br>(Autodock) | No of<br>Hydrogen<br>bond<br>interaction | Hydrogen - Aminoacid<br>interactions |
| --- | --- | --- | --- | --- |
| NCX1 | 2DPK | -2.67 | 5 | GLU A:452, THR A:415,<br>ASN A:417, |
| ATP2A1/SERCA1 | 6HEF | -4.09 | 3 | LEU A:797, ALA A:100 |
| CACNA1S | 2VAY | -3.49 | 8 | ARG A:74, LYS A :75,<br>GLU A:54, MET A:51,<br>LYS B:1522, GLU A:87,<br>GLU A:83 |
| S100 | 1A03 | -0.98 | 5 | LYS A:64, GLN A:66,<br>GLU A:67, ASN A:69 |
| Ca<sup>2+</sup>/calmodulin-<br>dependent protein kinase II | 2UX0 | +0.13 | 6 | TYR A:418, ARG A:502,<br>CYS A:518, SER A:498,<br>TYR A:516 |
| PKA C-α (Protein kinase A<br>catalytic subunit) | 2GU8 | -7.96 | 8 | VAL A:123, GLU A:127,<br>TYR A:330, THR A:51,<br>LYS A:72 |
| Triadin | 6QA5 | -3.19 | 5 | GLN A:431, ARG A:526,<br>GLY A:521 |
| BCL6 | 5N20 | +29.27 | 5 | ALA A:15, LEU A:19,<br>THR A:48 |
| PTEN | 1D5R | -2.02 | 4 | HIS A:93, GLN A:171,<br>CYS A:124, ARG A:130 |
| COX2 | 5F1A | -9.52 | 4 | ALA A:199, TYR A:385,<br>ASN A:382 |
| BCL2 | 1G5M | -4.65 | 8 | ASN A:182, THR A:7,<br>ARG A:6, ALA A:4, GLN<br>A:190, TRP A:195 |
| P21 | 5P21 | -4.57 | 8 | THR A:58, LYS A:16,<br>ASP A:38, PRO A:34,<br>ASP A:33, SER A:17,<br>GLU A:31 |
| CALRETICULIN | 3POW | -5.78 | 6 | GLU A:345, ASP A:63,<br>ASN A:329, LYS A:62,<br>GLN A:26 |

**Table 13:**
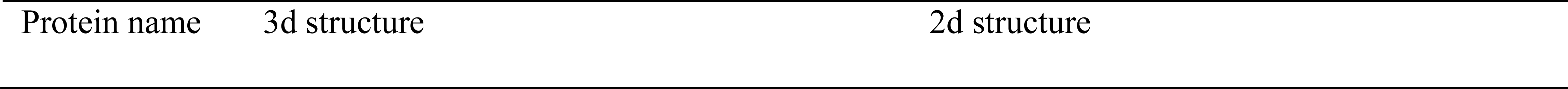
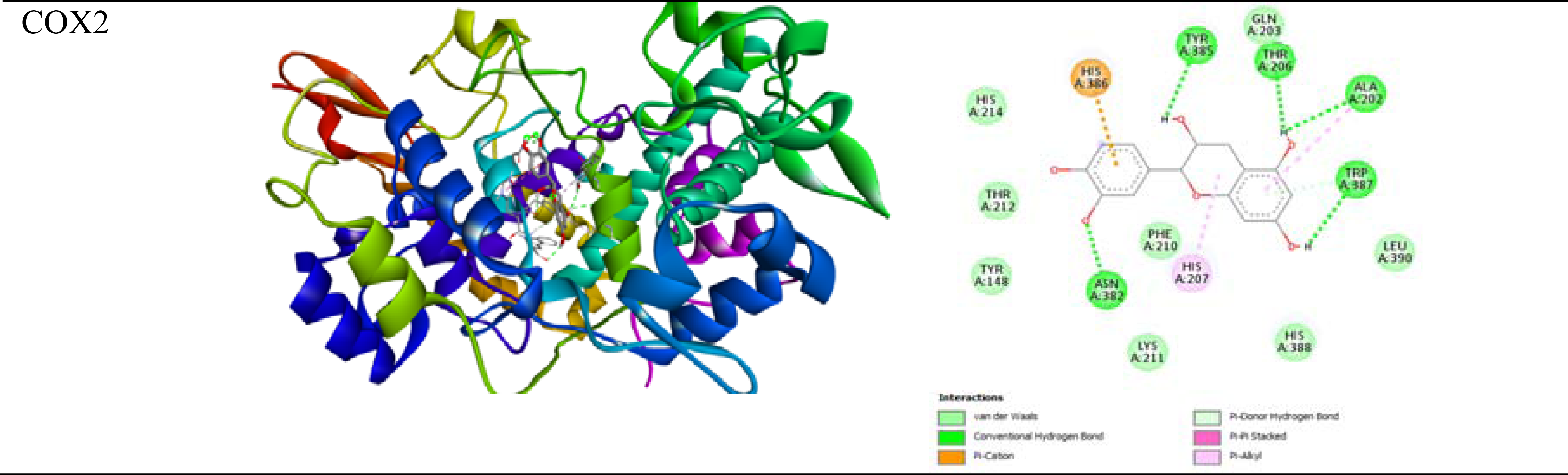
3D and 2D structure of Epicatechin with COX2.

**Table 14:**
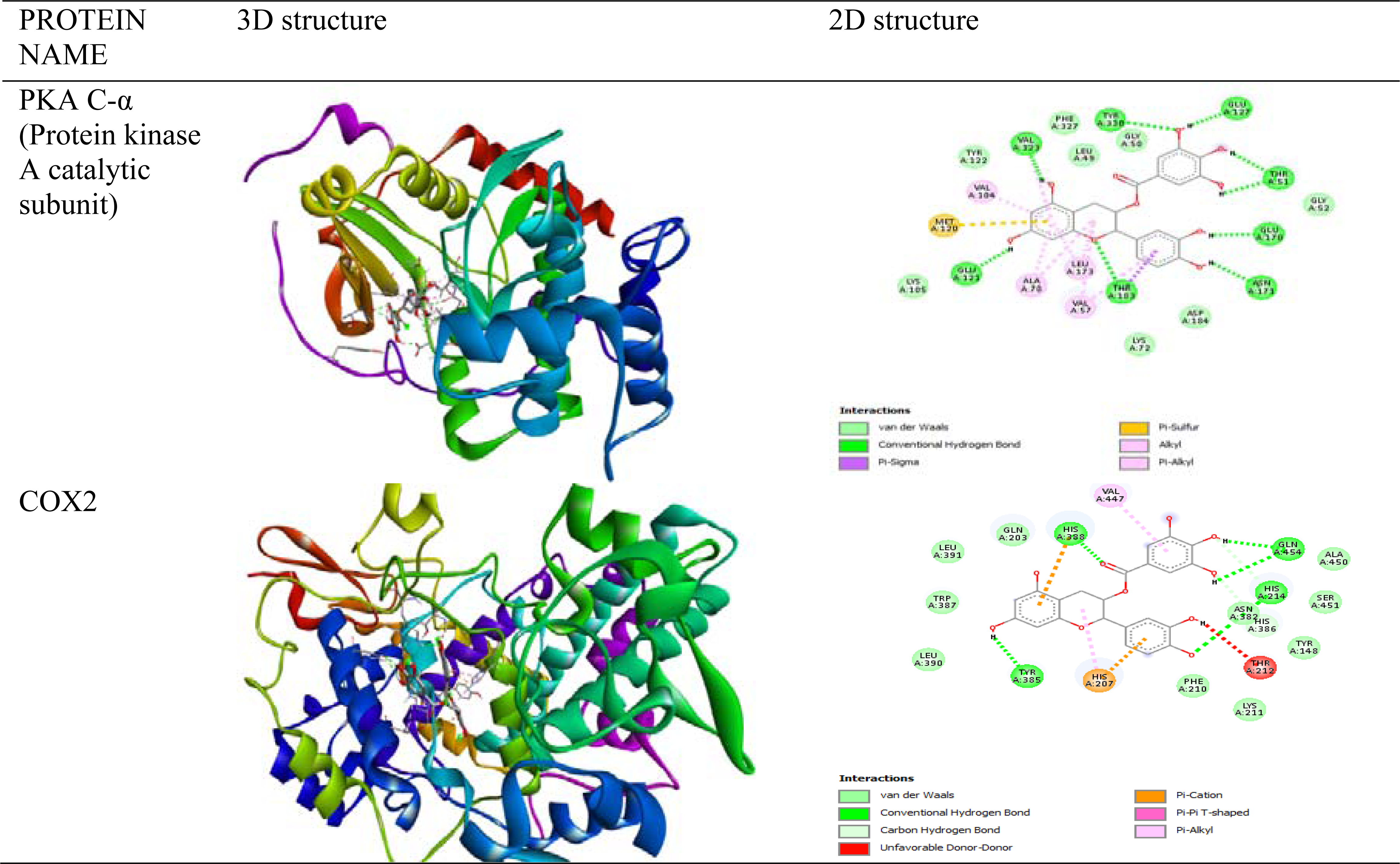
3D and 2D structure of Epicatechin Gallate with PKA C-α and COX2.

**Table 15:**
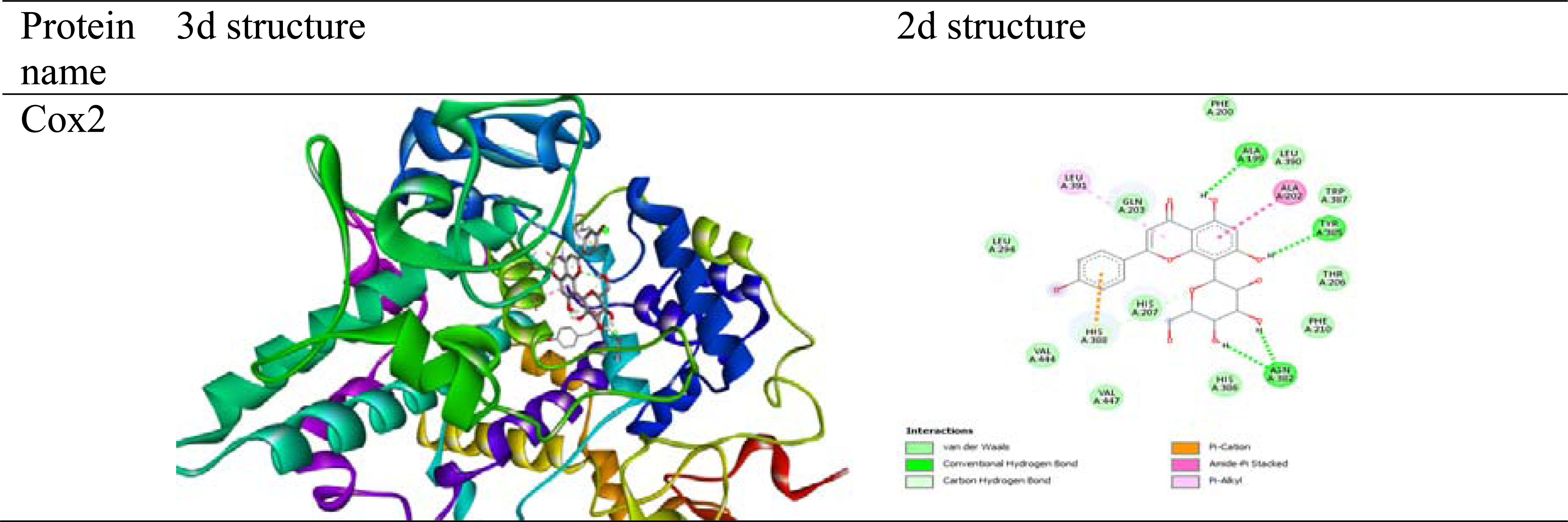
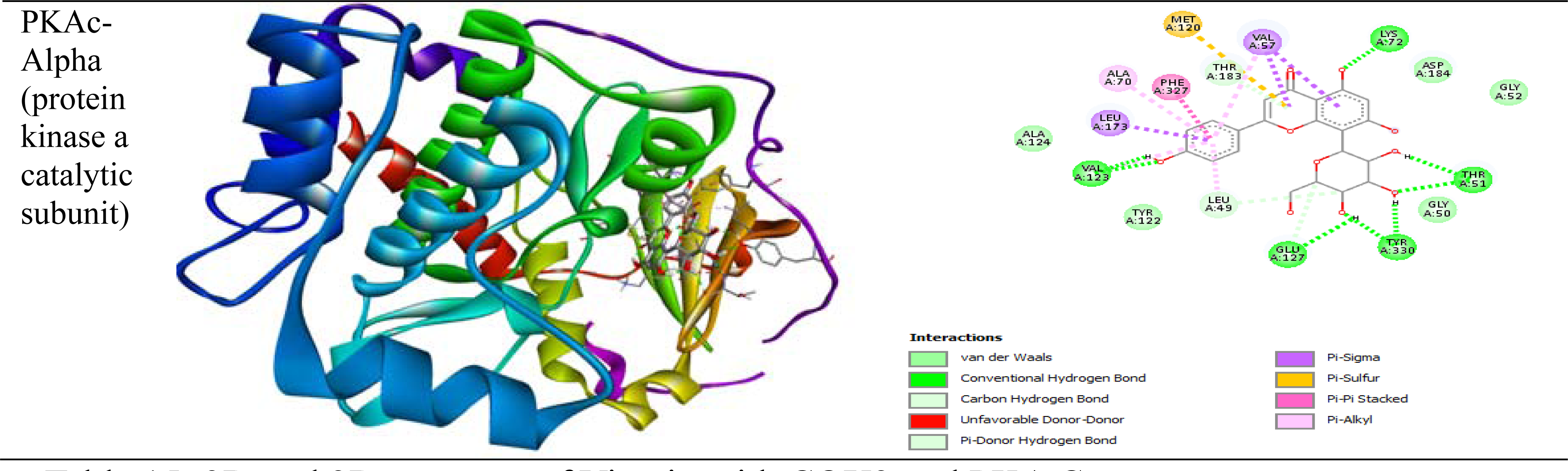
3D and 2D structure of Vitexin with COX2 and PKA C-α.

**Table 16:** A comprehensive binding energy results with hydrogen bonding interactions of molecular docking of Epicatechin with Proteins embroiled in the signaling of Ca2+ overloading through maestro.

| PROTEIN NAME | PDB ID | Glide Energy | Docking score (Schrodinger) | No. Hydrogen bond interaction | Interacting residues |
| --- | --- | --- | --- | --- | --- |
| NCX1 | 2DPK | -32.6401 | -4.721 | 3 | GLU 454, ASP 421, ASP 447 |
| ATP2A1/SERCA1 | 6HEF | -30.0589 | -5.262 | 1 | SER 936 |
| CACNA1S | 2VAY | -36.9006 | -5.515 | 5 | GLU A:54, ASP A:78, LYS B:1522 |
| S100 | 1A03 | -28.7924 | -4.548 | 3 | GLU A:67, GLN A:66 |
| Ca<sup>2+</sup>/calmodulin-dependent protein kinase II | 2UX0 | -30.8013 | -4.285 | 5 | GLU 430, GLU 432, ARG 482, ARG 502, GLY 1521 |
| PKA C-α (Protein kinase A catalytic subunit) | 2GU8 | -46.8886 | -6.508 | 3 | THR A:51, THR A:183 |
| Triadin | 6QA5 | -31.5939 | -4.875 | 4 | ASP 420, ASP 450, ASP 524, ARG 526 |
| BCL6 | 5N20 | -25.5627 | -4.519 | 2 | ALA 15, SER 93 |
| PTEN | 1D5R | -32.2152 | -4.491 | 3 | GLN 171, ARG 130, ASP 92 |
| COX2 | 5F1A | -43.4256 | -7.641 | 4 | THR 212, HIE 214, HIS 207, TYR 385 |
| BCL2 | 1G5M | -39.5288 | -6.156 | 3 | ALA 4, THR 7, TYR 9 |
| P21 | 5P21 | -46.77 | -5.590 | 4 | THR 58, SER 17, GLY 60, GLU 31 |
| CALRETICULIN | 3POW | -41.8368 | -6.475 | 4 | ASN 329, ASP 63, GLN 26, GLU 345 |

**Table 17:** A comprehensive binding energy results with hydrogen bonding interactions of molecular docking of Epicatechin Gallate with Proteins embroiled in the signaling of Ca2+ overloading through maestro.

| PROTEIN NAME | PDB ID | Glide energy | Docking score (Schrodinger) | No of Hydrogen bond interaction | Hydrogen - Aminoacid interactions |
| --- | --- | --- | --- | --- | --- |
| NCX1 | 2DPK | -46.7078 | -5.249 | 3 | ASP 447, ASP 421, ,<br>GLU 454 |
| ATP2A1/SERCA1 | 6HEF | -43.2664 | -5.657 | 3 | TRP 107, SER 936,<br>ASN 111 |
| CACNA1S | 2VAY | -42.6684 | -1.788 | 5 | GLU A: 54, ASA<br>A:78 |
| S100 | 1A03 | -35.6653 | -3.212 | 5 | GLN A:66, GLU<br>A:67, GLN A:71,<br>ALA B:87 |
| Ca<sup>2+</sup>/calmodulin-<br>dependent protein kinase II | 2UX0 | -39.0745 | -3.951 | 4 | ARG 502, ARG 482,<br>GLU 432 |
| PKA C-α (Protein kinase A<br>catalytic subunit) | 2GU8 | -63.8962 | -8.311 | 5 | ASN C:520, ARG<br>C:519, THR A:183,<br>ASP A:184, THR<br>A:51 |
| Triadin | 6QA5 | -37.6003 | -4.245 | 3 | ASP 524, ASP 459,<br>HIS 493 |
| BCL6 | 5N20 | -37.4316 | -3.820 | 1 | ALA 52 |
| PTEN | 1D5R | -52.5515 | -5.051 | 5 | ASP 162, TYR 16,<br>ASP 92 |
| COX2 | 5F1A | -53.645 | -7.402 | 4 | HIS 207, ALA 199,<br>TYR 385 |
| BCL2 | 1G5M | -46.7951 | -5.703 | 3 | THR 7, GLY 5, GLN<br>190 |
| P21 | 5P21 | -55.0867 | -5.463 | 5 | LYS 16, SER 17, GLY<br>60, THR 58, ASP 30 |
| CALRETICULIN | 3POW | -47.0308 | -5.296 | 6 | GLN 26, LYS 62, ASP<br>63, ASN 329, GLU<br>345 |

**Table 18:**
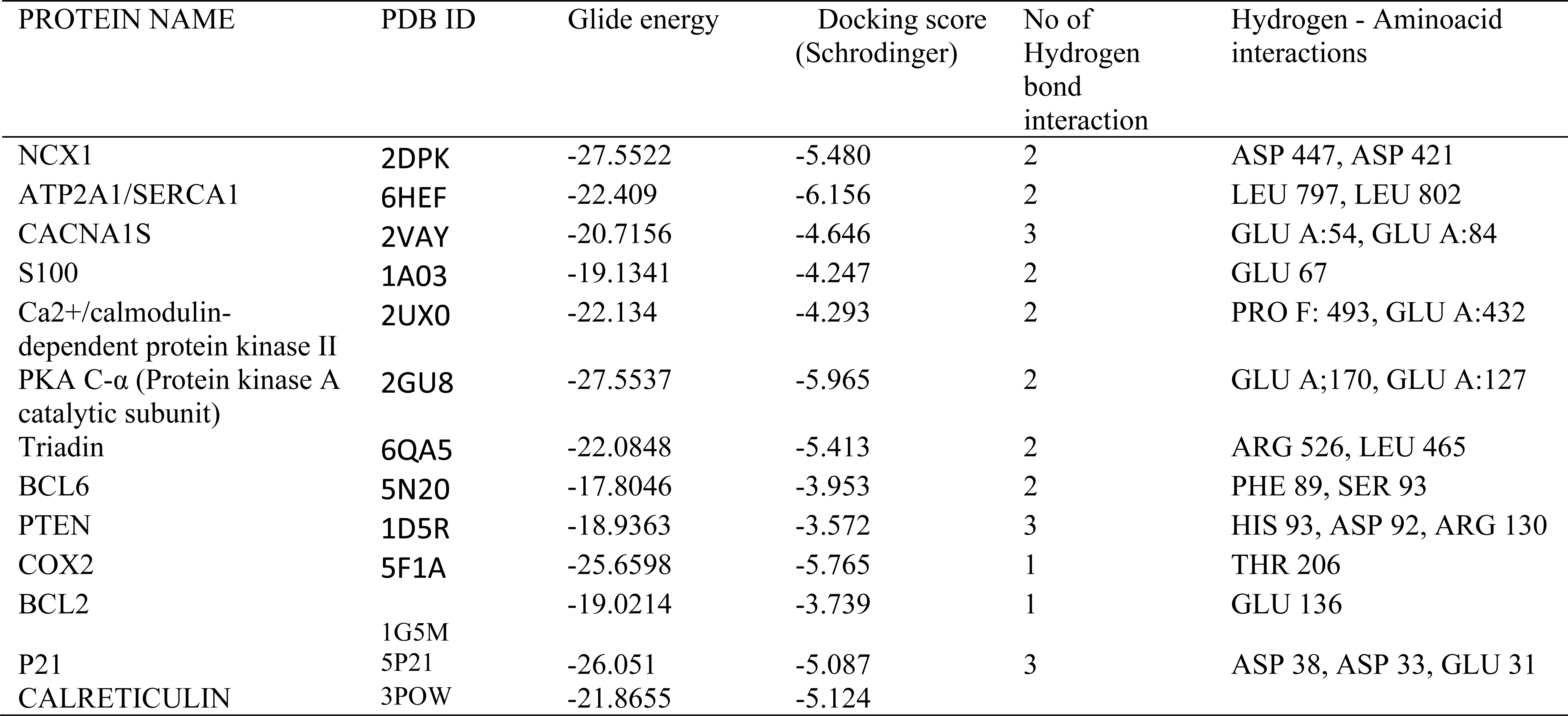
A comprehensive binding energy results with hydrogen bonding interactions of molecular docking of Tyramine with Proteins embroiled in the signaling of Ca2+ overloading through maestro.

**Table 19:**
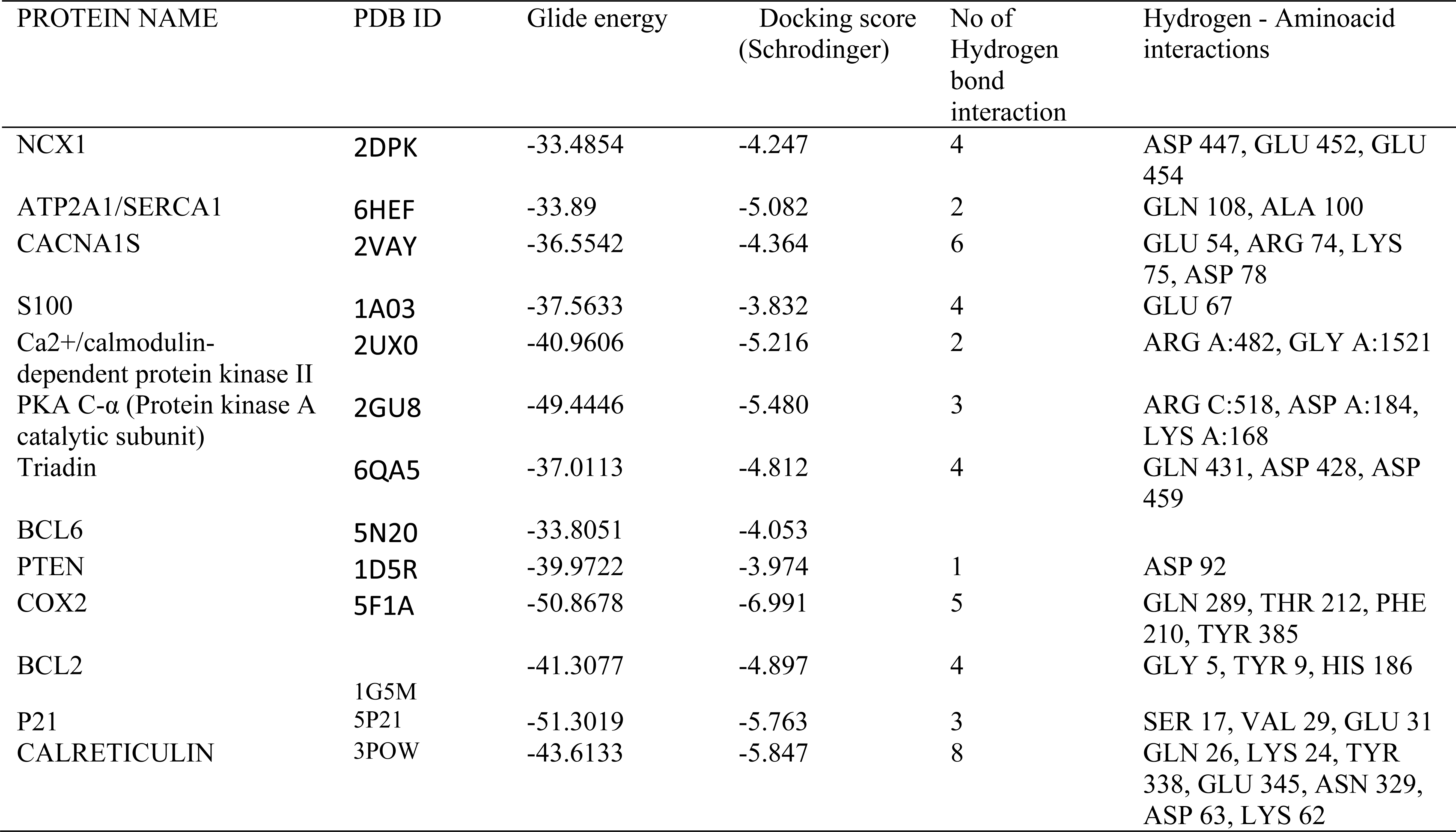
A comprehensive binding energy results with hydrogen bonding interactions of molecular docking of Vitexin with Proteins embroiled in the signaling of Ca2+ overloading through maestro.

During the process of modern and advance drug development studies, Molecular docking and molecular dynamics simulation studies have become fundamental [36]. Molecular docking is the process used to determine optimal binding sites between a ligand to a protein in light of preferred orientation of several parameters. In the pharmaceutical sector comparative protein-ligand docking is utilized to assess efficaciously evaluate the accuracy of molecular interactions and identify prospective drug candidates with enhanced precision and efficiency [37]. The utilization of advanced algorithmic software platforms, such as Auto-dock, Auto-dock vina, and Maestro are prominently deployed in this context. These software packages employ distinct scoring functions that facilitate the computation of binding affinity and binding energy, yielding crucial insights into the nature of molecular interactions to identify viable novel drugs candidates [38]. Furthermore, comparative molecular docking can be used to identify potential drug candidates for a variety of diseases including cardiovascular disease. By using this approach, it is possible to compare the results of multiple interactions obtained from auto-dock, auto-dock vina and glide can lead to the identification of drugs which will modulate multiple targets, resulting in broader therapeutic effects. Hence, comparative molecular docking is a very most valuable in drug discovery, as it enables the efficient identification of potential drug candidates and provides valuable insights into the molecular interactions between proteins and ligands. By utilizing advanced software and scoring functions, we can identify and optimize drug candidates for a variety of diseases, ultimately leading to the development of more effective drugs. As a consequence, Glide (Maestro), Autodock/ Autodock vina was compared to perceive the most reliable docking routines to track down the potent inhibitors from COC against thirteen proteins. Docking score of four potential secondary compounds from COC with thirteen proteins were indicated in the above tables. As contrasted with whole proteins and ligands, epicatechin gallate with PKA Cα experienced high negative docking scores in Glide (Maestro), Autodock and Autodock Vina.

In cardiac muscle cells, Protein kinase C alpha (PKCα) plays a role in Ca2+ handling and regulating cardiac contractility through the inflection of its activity cause dephosphorylation on pump inhibitory protein phospholamban (PLB), a pump inhibitory protein of sarcoplasmic reticulum Ca2+ ATPase-2 (SERCA-2) and alters Ca2+ transients in the sarcoplasmic reticulum [39]. PKCα also regulates mitochondrial aconitase activity through PKC dependent phosphorylation which may influence tricarboxylic acid cycle activity in diabetic hearts and impair the function of mitochondria [40]. According to the recent study, PKCα inhibition has reduce left ventricular fibrosis in myocardial infarction [41]. The results of protein-ligand docking studies suggest that PKCα interacts favourably with epicatechin gallate, a secondary metabolite of COC.

Utilizing Desmond, 2000ns of molecular dynamics simulation were performed to investigate the molecular flexibility and structural behaviour of PKA Cα with the epicatechin gallate found in COC. Initially, the protein-ligand complex was bound between 328ns and 1286ns with Protein RMSD values ranging from 1.87Å to 2.27Å and Ligand RMSD values ranging from 3.97Å to 4.63Å. However, between 1287ns and 1668ns, the complex showed instability with Protein RMSD values increasing to 2.85Å and Ligand RMSD values reaching 4.67Å. Subsequently, from 1668ns to the end of the simulation at 2000ns, the protein-ligand complex stabilized with Protein RMSD values ranging from 2.78Å to 3.34Å and Ligand RMSD values ranging from 4.91Å to 5.82Å. As a result, a stable structure with an RMSD value of approximately 5.84Å was observed during the simulation, which was superior to the reference RMSD value when PKA Cα was simulated with epicatechin gallate for a duration of 2000ns.

The Root Means Square Fluctuations (RMSF) is to depict the associations between the epicatechin gallate and local fluctuations of the PKAC-alpha. From the figure (3), RMSF values of the epicatechin gallate atoms were significantly higher than those of the PKAC-alpha residues indicating that the epicatechin gallate exhibits higher flexibility and undergoes more pronounced structural changes than the protein during the simulation time. In contrast, the binding site and its surrounding residues showed relatively low levels of fluctuation with RMSF values ranging from 1.59 Å to 2.42 Å which indicates that the epicatechin gallate binding stabilizes the local protein structure and reduces its conformational flexibility. Accordingly, the plot shows the reliability and fluctuations of PKA Cα as well as the specific amino acid’s interaction with epicatechin gallate.

**Figure 3:**
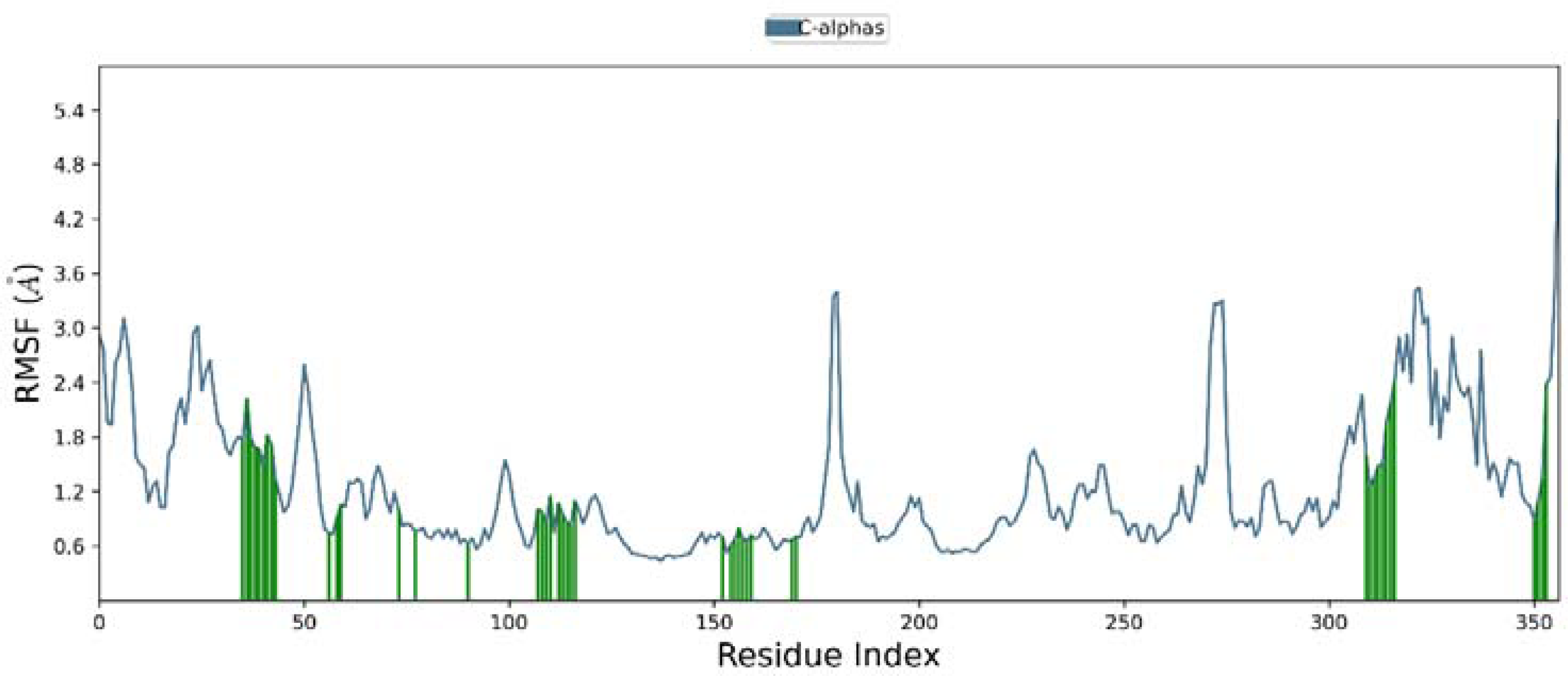
Root means square fluctuations (RMSF) plot for PKA C-α – Epicatechin gallate during 2000ns of molecular dynamics.Waveforms of PKA C-α is shown in the blue colour and the green line portrays epicatechin gallate interactions with amino acids of PKA C-α.

In the consistent areas of the PKA Cα with epicatechin gallate, figure (4) revealed hydrogen bonds, water bridges, and hydrophobic interactions. From figure (4), LEU 49 formed hydrogen bond interactions at a frequency of 0.329 and hydrophobic interactions at a frequency of 0.025, while also forming water bridges at a frequency of 0.142. In addition to LEU 49, other amino acid residues also formed stable contacts with the ligand. For instance, THR51 formed hydrogen bond interactions with a frequency of 0.495 and water bridges with a frequency of 0.183. GLU121 formed hydrogen bond interactions with a frequency of 0.552 and water bridges with a frequency of 0.163. VAL123 formed hydrogen bond interactions with a frequency of 0.612 and water bridges with a frequency of 0.108. ASP184 formed hydrogen bond interactions with a frequency of 0.955 and water bridges with a frequency of 0.113. Finally, ASP328 formed hydrogen bond interactions with a frequency of 0.609 and water bridges with a frequency of 0.422. Among that, residue ASP184 maintained the specific interaction for 90% of the simulation time, as indicated by its fraction contact value of 0.955. Similarly, residue ASP328 maintained the specific interaction for 60% of the simulation time, as indicated by its fraction contact value of 0.609. Residue VAL123 maintained the specific interaction for 60% of the simulation time, as indicated by its fraction contact value of 0.612. Finally, residue ASP184 maintained the specific interaction for 90% of the simulation time, as indicated by its fraction contact value of 0.955. As the result of the interaction frequencies indicates that specific interactions are maintained over the 2000ns which provides valuable insights into the binding mechanism. As a subsequence of this graph, the simulation of PKA Cα with epicatechin gallate was determined that there was a number of hydrogen bonds with a decent frequencyand have the potential for rational drug design.

**Figure 4:**
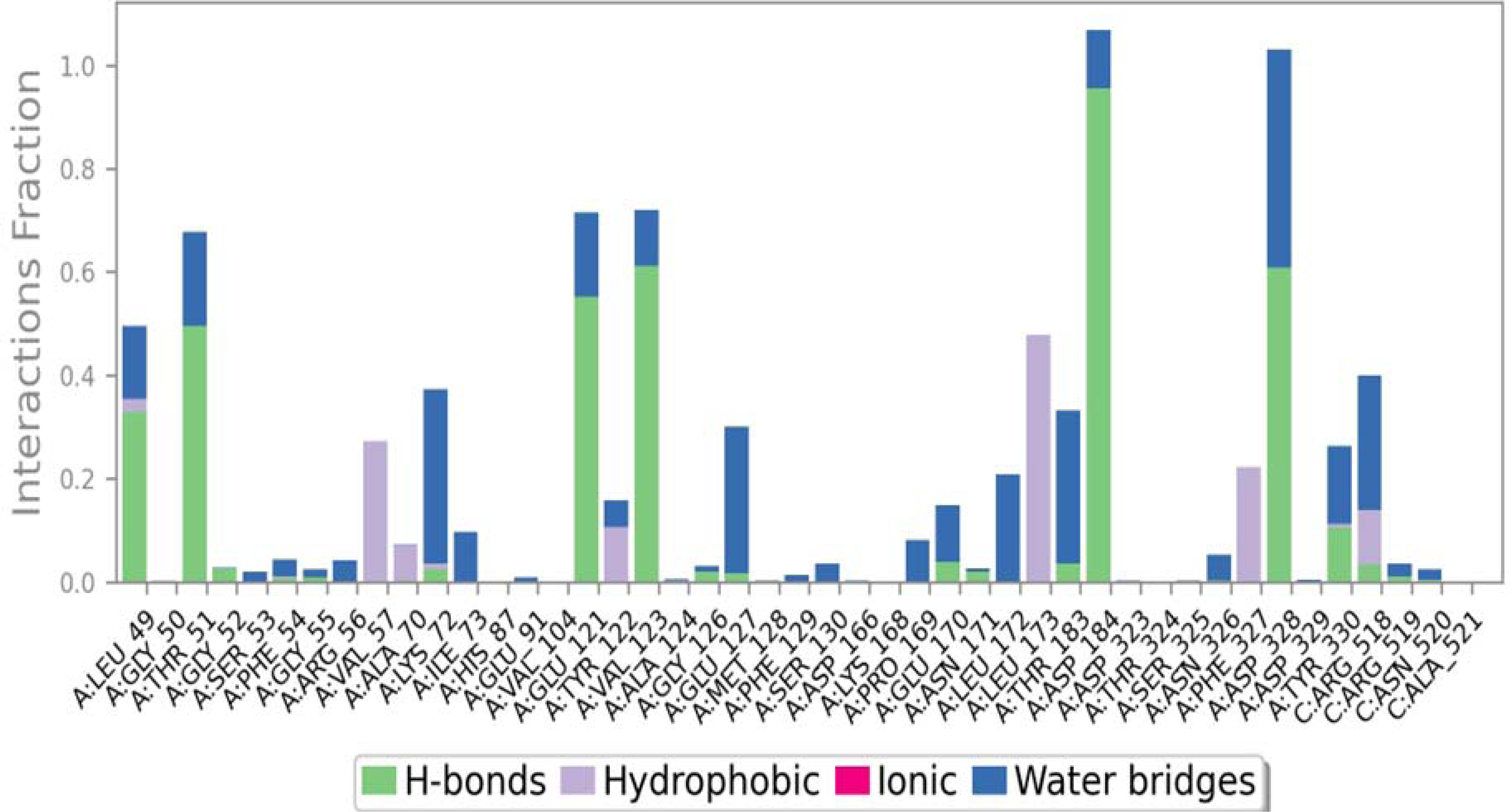
The plot represents the hydrogen bonding interactions of PKA C-α – Epicatechin gallate during 2000 ns MD simulation.

From the beneath heatmap figure (5), residues made contacts based on the time on the x-axis and the residues alluded to the y-axis for making contact. Some residues have multiple specific contact with in each trajectory frame as represent in darker orange colour, as shown by the scale to the right of the plot, while others represent in different shades. During most interactions, ASP184, ASP328, VAL123, THR51 and GLU121 typically made direct hydrogen bond interactions with the ligand till 2000th ns. The darker orange shade demonstrates that ASP328, VAL123 and ASP184 could make multiple contacts simultaneously. Accordingly, the number of specific contacts that the PKA Cα has with the epicatechin gallate throughout the trajectory is efficaciously shown in below heatmap.

**Figure 5:**
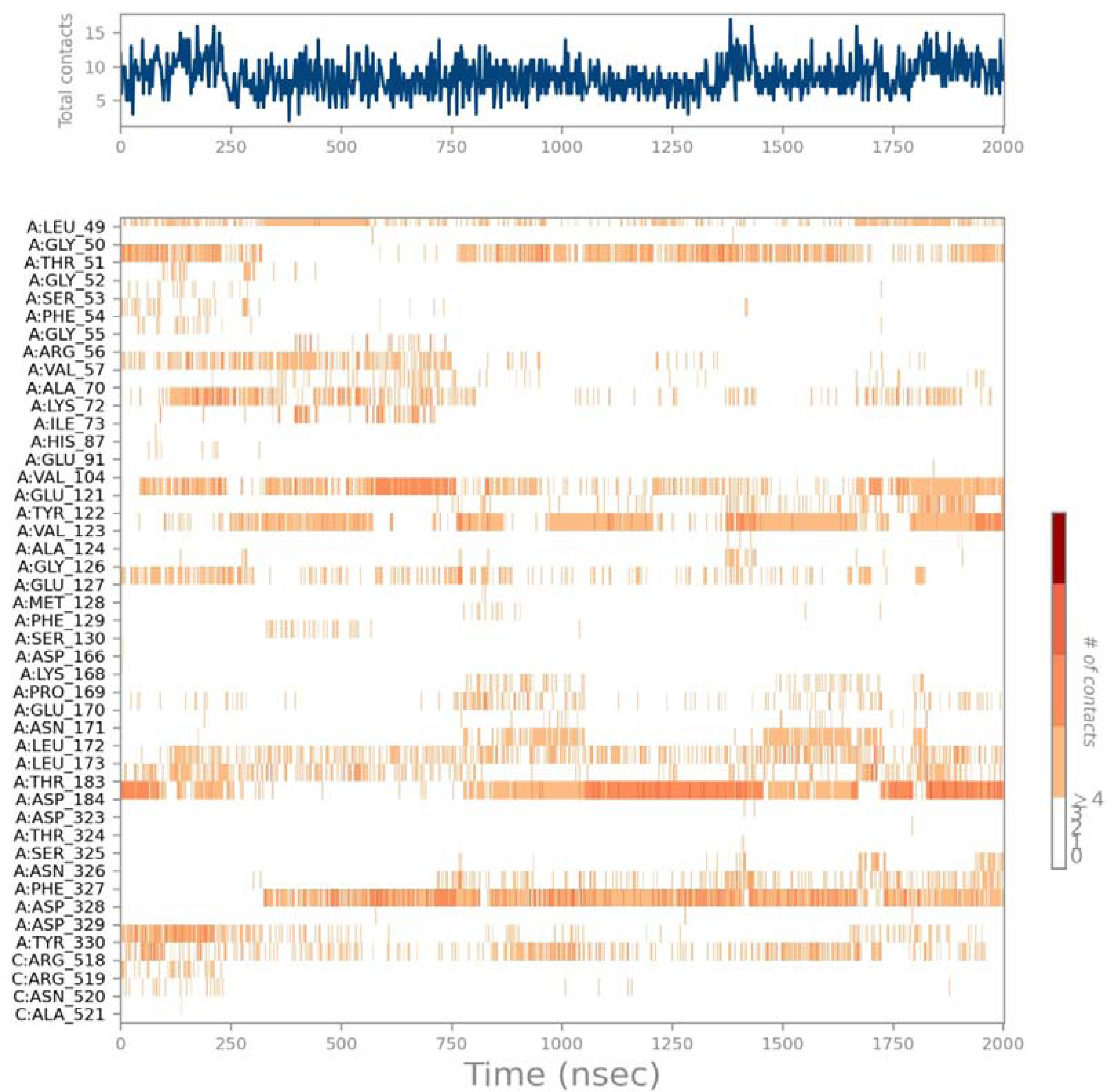
A heat map depicting the complete number of interactions formed by the PKA Cα to the Epicatechin gallate over 2000ns. White signifies zero, pine signifies one contact, light orange shade signifies two contacts, dark orange shade signifies three contacts and red shade signifies greater than four contacts.

From the figure (6), A dial plot and bar plots of the same colour accompany this plot, which provides a summary of the conformational evolution of all rotational bond of epicatechin gallate with individual rotatable bond torsion. Besides, each conformation from the dial plot commences in the midpoint of the radical plot as well as the time assessment is charted radially outwards. In addition, Summing the potentials of the related torsions of the Epicatechin gallate reveals potential torsional information, akin to that bar plots depict the probability density of the torsions. Based on the analysis of the histogram and torsion potential relationship, several rotational bonds of epicatechin gallate can capable to undergo and maintain a protein bound conformation.

**Figure 6:** An interactive torsion plot of epicatechin gallate with two-dimensional schematics of coloured rotatable bonds throughout the ligand simulation. Y-axis indicates the values of the potential in kcal/mol.

**Figure 7:**
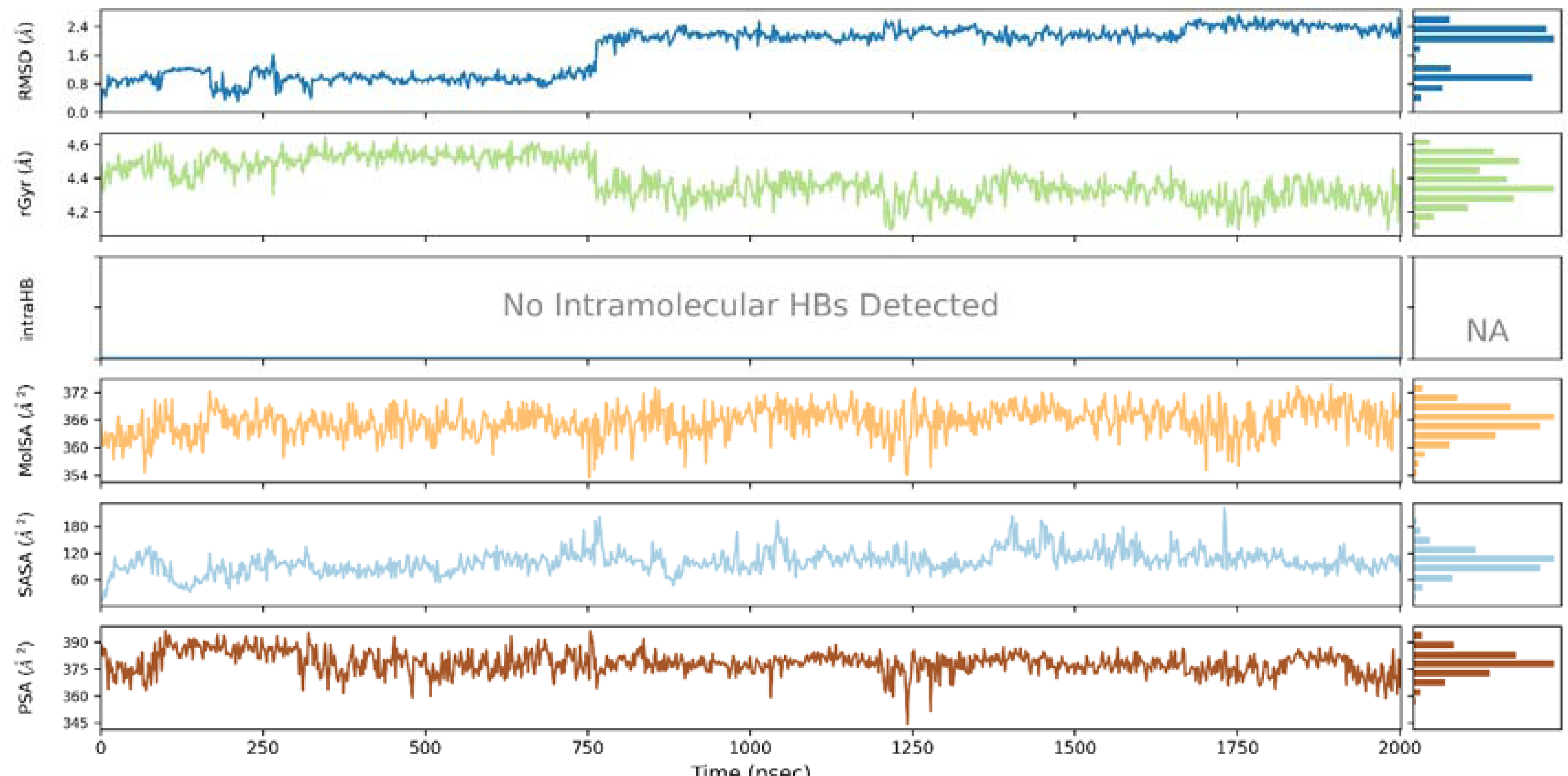
The plot and bar charts of Epicatechin gallate properties (i.e.) Ligand RMSD, Radius of Gyration (rGyr), Intramolecular Hydrogen Bonds (intraHB), Molecular Surface Area (MolSA), Solvent Accessible Surface Area (SASA) and Polar Surface Area (PSA) during 2000 ns MD simulation. x-axis indicates property value and y-axis indicates simulation time.

**Figure 8:**
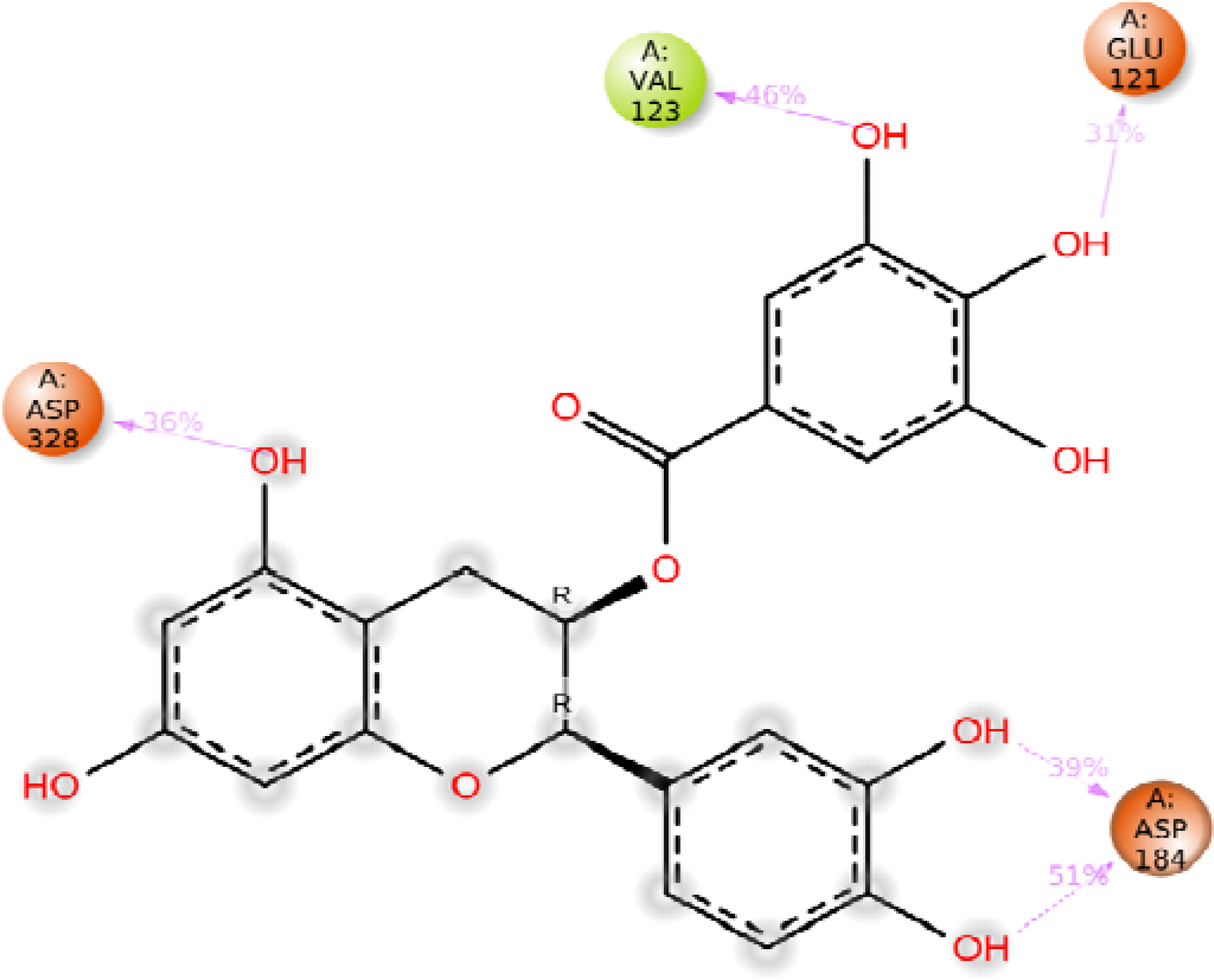
A detailed 2D schematic of the interactions between the atoms of Epicatechin Gallate and the neighbouring residues of a PKA Cα.

Based on the analysis of Epicatechin gallate properties was investigated by utilizing the following parameters: Ligand RMSD, Radius of Gyration (rGyr), Intramolecular Hydrogen Bonds (intraHB), Molecular Surface Area (MolSA), Solvent Accessible Surface Area (SASA) and Polar Surface Area (PSA) during the 2000 ns of MD simulation. Epicatechin gallate showed a substantial parameter in rGyr, MolSA, SASA, PSA and there was no intraHB.

Overall, five hydrogen bond interactions were identified with the residue ASP328 formed hydrogen bond interactions for 36% of the MD trajectory, residue VAL123 for 46% of the MD trajectory, residue GLU121 for 31% of the MD trajectory, and residue ASP184 for 51% of the MD trajectory. Additionally, residue ASP184 established two strong hydrogen bonds with the OH anions of epicatechin gallate for 39% of the MD trajectory, indicating adequate interactions.

Henceforth, 2000ns MD simulations indicate that epicatechin gallate has sufficient conformational flexibility. According to the above result, VAL123, ASP184, GLU121 and ASP328 was crucial for stabilizing epicatechin gallate within PKA Cα (2GU8) binding pocket and therefore epicatechin gallate is consequently stable in catalytic pocket. By the activation of PKA Cα, several proteins in cardiomyocytes are phosphorylated, resulting in an increase in contraction strength and frequency. Supplementally, when PKA Cα regulates PLB phosphorylation, the endoplasmic reticulum’s Ca2+ uptake level elevates [42].For that reason, activation of the regulatory enzyme PKA Cα have high possibilities leads to felicitous phosphorylation resulting positive inotropy and lusitropy. As an effective cardio tonic, COC(hawthorn) utilized to treat various cardiac diseases [43]. During myocardial infarction, epicatechin gallate from COC is capable of inhibiting caspase3. In antecedent research from my lab epicatechin gallate shows efficacious against caspase3[44]. As a consequence, epicatechin gallate has high possibilities to inhibit PKA Cα for felicitous phosphorylation.

## Conclusion

Overburden of Ca2+ mediates a significant contributor for Heart failure, Atrial failure and arrhythmias. According to the intensive assessment of protein - protein interaction, uncovers various proteins are enrolling in numerous activities of Ca2+ overloading. In present study, molecular docking analysis distinguished puissant secondary metabolite - Epicatechin gallate from COC with harmless ADMET profile, high affinities and potential inhibition towards abnormalities inducing proteins cardio Ca2+. Besides, the 2000ns of molecular dynamics simulation studies affirmed their conformational rigidity nature and found the epicatechin gallate have a substantial binding mode. The current insillico research suggests that epicatechin gallate from COC may inhibit cardiac Ca2+ abnormalities. Furthermore, in light of current insillico data, the experimental investigation can aid to the advancement of novel medication to combat myocardial infarction complications

## Acknowledgements

“We would like to express our sincere gratitude to RUSA-2.0 Biological Sciences for their financial and software support and to USIC-BDU for allowing us access to their computer facilities. We would also like to extend our appreciation to senior scientist Vinod Devaraji at Schrodinger for providing technical support to our Molecular Dynamics studies.”

